# Acute AMPK activation does not adequately stimulate insulin signaling in skeletal muscle models of Myotonic Dystrophy Type 1

**DOI:** 10.1101/2025.11.04.686135

**Authors:** Ofosu Adjei-Afriyie, Sally Spendiff, Ricardo Carmona-Martinez, Alexander Manta, Andreas Hentschel, Kelly Ho, Daniel O’Neil, Leanne Dawe, Andreas Roos, Vladimir Ljubicic, Alexander MacKenzie, Aymeric Ravel-Chapuis, Hanns Lochmüller

**Affiliations:** Department of Cellular and Molecular Medicine, University of Ottawa, 451 Smyth Road, Ottawa, Ontario, Canada, K18 HM5; Children’s Hospital of Eastern Ontario Research Institute, 401 Smyth Road, Ottawa, Ontario, Canada, K1H 8L1; The Eric Poulin Centre for Neuromuscular Disease, University of Ottawa, 451 Smyth Road, Ottawa, Ontario, Canada, K18 HM5; Department of Kinesiology, McMaster University, 1280 Main Street West, Hamilton, Ontario, L8S 4K1; Department of Medicine, University of Toronto, 1 King’s College Circle, Toronto, Ontario, Canada, M5S 1A8; Leibniz-Institute for Analytical Science -ISAS-e.V., Otto-Hahn-Strasse 6b, 44227 Dortmund, Germany; Department of Pediatric Neurology, Centre for Neuromuscular Disorders, University Duisburg-Essen, 45147 Essen, Germany; Department of Neurology, Medical Faculty and University Hospital Düsseldorf, Heinrich Heine University, 40225 Düsseldorf, Germany; School of Pharmaceutical Sciences, Univeristy of Ottawa, 451 Smyth Road, Ottawa, Ontario, Canada, K18 HM5; Brain and Mind Research Institute, University of Ottawa, 451 Smyth Road, Ottawa, Ontario, Canada, K18 HM5; Department of Neuropediatrics and Muscle Disorders, Medical Center – University of Freiburg, Faculty of Medicine, Mathildenstr. 1, 79160 Freiburg, Germany; Centro Nacional de Análisis Genómico, Baldiri Reixac 4, 08028 Barcelona, Spain

**Author notes:** Children’s Hospital of Eastern Ontario Research Institute, 401 Smyth Road, Ottawa, Ontario, Canada. To whom correspondence should be addressed: Hanns Lochmüller.

**Keywords:** Myotonic dystrophy type 1 (DM1), exercise, insulin, AMPK, PGC-1α, Akt, AS160, GLUT4

## Abstract

Myotonic Dystrophy Type 1 (DM1) is a multisystemic neuromuscular disorder characterized by skeletal muscle weakness, muscle atrophy, myotonia, cognitive impairments, gastrointestinal complications, and insulin resistance. While insulin resistance is well characterized in type 2 diabetes, its pathomechanism in DM1 remains unclear. Our study aims to elucidate the pathomechanism of insulin resistance in DM1 and how the pathway responds to AMPK stimulation. Proteomic analysis from sedentary wildtype and sedentary HSA-LR mice, a common DM1 mouse model, revealed downregulation of the AMPK-PGC-1α axis. Analysis of sedentary HSA-LR mice and exercised HSA-LR mice revealed activation of the AMPK-PGC-1α axis in exercised animals. To investigate this pathway, we treated WT and HSA-LR mice with the AMPK activator AICAR to examine the impact of AMPK stimulation on insulin signaling in DM1. This revealed impaired responses in the insulin pathway activation in the HSA-LR mice. Next, we examined whether these differences extended to a human model by treating control and DM1 myotubes with insulin and/or AICAR. In DM1 myotubes, both treatments produced dampened responses of key insulin signaling intermediates compared to controls. Taken together, these results suggest impaired activation of insulin signaling pathways in DM1 models and confirm the presence of insulin resistance with an impaired response to acute AMPK stimulation.

---

1. **Introduction**

Myotonic dystrophy 1 (DM1), also known as Steinert’s Disease, is a multisystemic neuromuscular disease that affects approximately 1 in 8500 people worldwide, making it the most common adult-onset muscular disease (Chau and Kalsotra, 2015; Yotova et al., 2005). Due to a founder effect, there is a strikingly high prevalence of 1 in 550 affected individuals in the Charlevoix and Saguenay-Lac-Saint-Jean regions of Quebec, Canada (Chau and Kalsotra, 2015; Yotova et al., 2005).

DM1 patients present with a heterogeneous set of clinical symptoms, including myotonia, characterized as prolonged muscle contraction. Other symptoms include progressive muscle weakness, muscle atrophy, and metabolic abnormalities, such as insulin resistance (Nieuwenhuis et al., 2019; Mateus et al., 2021). Insulin signaling is integral to the energy status of the cell, governing the intake of glucose to fuel energy demanding processes like cell growth and division. In DM1, systemic glucose disposal is diminished by 15-25%, with up to a 70% decrease in insulin sensitivity observed in the forearm muscle of DM1 patients (Moxley et al., 1977, 1984). When insulin signaling is impaired in DM1, it can exacerbate coexisting symptoms such as muscle weakness and atrophy (Nieuwenhuis et al., 2019). Additionally, it can cause significant impairments to skeletal muscle, cardiac, adipose and brain tissue in DM1 (Nieuwenhuis et al., 2019).

DM1 arises from a CUG-expansion repeat in the 3’ untranslated region of the dystrophia myotonica protein kinase (*DMPK*) gene (Brook, 1992; Caskey et al., 1992; Mahadevan et al., 1992). When transcribed, these repeats cause a stable hairpin loop of RNA which sequesters Muscleblind-like 1 (MBNL1), an RNA-binding protein, to the nucleus, resulting in its subsequent loss of function (Chau and Kalsotra, 2015). In parallel, CUG-binding protein 1 and Elav-like family member 1 (CELF1), an RNA-binding protein that functions antagonistically to MBNL1, is hyperactivated in DM1 skeletal muscle cells and contributes to mis-splicing events in pre-mRNA (Philips et al., 1998; Kuyumcu-Martinez et al., 2007; Chau and Kalsotra, 2015). CELF1 aberrantly excludes exon 11 in the insulin receptor (*INSR*) gene which contributes to insulin resistance in patients (Renna et al., 2017, 2019; Savkur et al., 2001). Insulin resistance in DM1 reflects dysregulation of downstream insulin receptor effectors that govern glucose uptake, cell metabolism and muscle trophism in DM1 (Renna et al., 2017, 2019).

The insulin signaling cascade contributes to cellular growth, development, and differentiation (Boucher et al., 2014; Renna et al., 2019). When the insulin hormone binds to the insulin receptor, the receptor phosphorylates and activates a signaling cascade that causes glucose transporter 4 (GLUT4) translocation to the plasma membrane, which allows the influx of glucose into the cell (Figure 1A) (Zerial and McBride, 2001). Once the insulin receptor is phosphorylated, a cascade of signaling events leads to the phosphorylation of Akt. Subsequently, Akt phosphorylates Akt substrate of 160 kDa (AS160), a GTPase-activating protein (GAP) for Rab proteins (Mîinea et al., 2005). In its native state, when AS160 interacts with Rab proteins, mobilization of GLUT4 storage vesicles (GSV) to the plasma membrane is inhibited. However, when Akt phosphorylates AS160 on threonine-642, it inhibits its GAP activity and promotes a greater proportion of active GTP-bound Rabs and increases GSV translocation to the plasma membrane (Figure 1A) (Mîinea et al., 2005).

**Figure 1.**
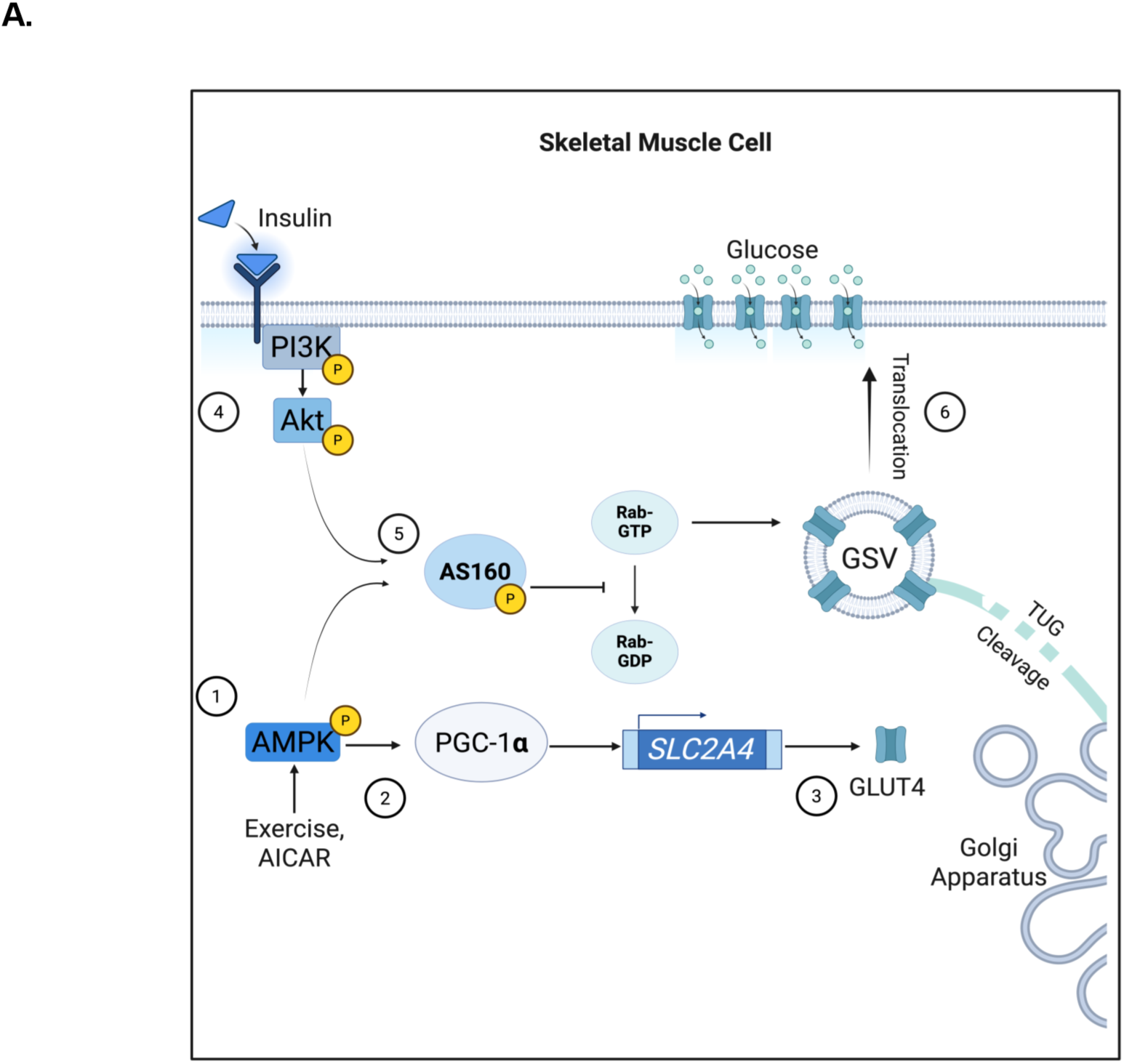
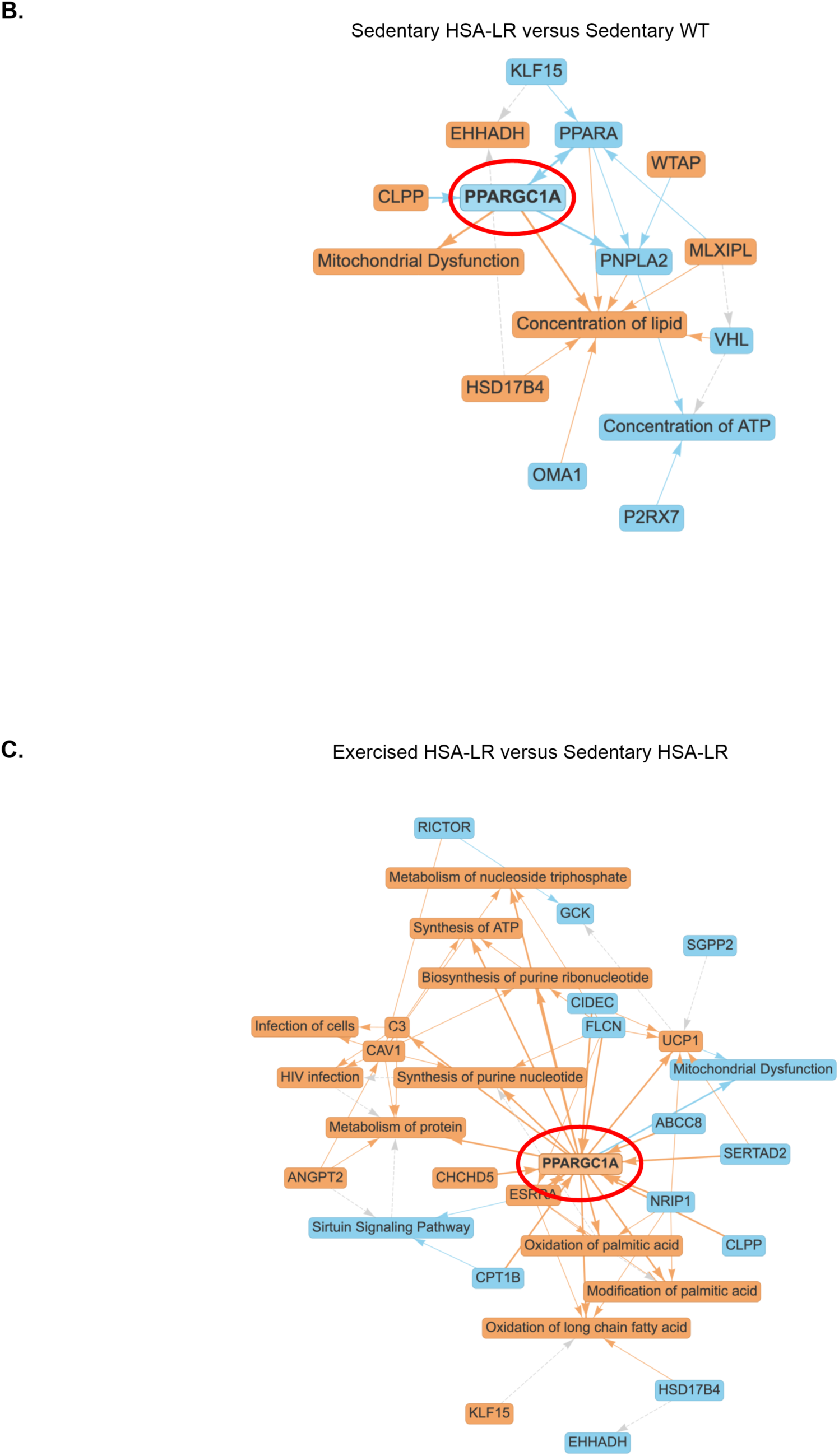
Predicted interactions of PGC-1α and insulin signaling pathway. **A.** Graphical depiction of proposed interactions between insulin signaling and exercise pathways in skeletal muscle. Adenosine monophosphate activated protein kinase (AMPK) stimulation via exercise or AICAR (1) can drive peroxisome proliferator-activated receptor gamma coactivator 1 alpha (PGC-1α) activity and expression (2) to increase expression of glucose transporter 4 (GLUT4) (3). Insulin binds to the insulin receptor which activates Akt (4) which leads to Akt substrate of 160 (AS160) stimulation (5). AS160 stimulation then drives GLUT4 storage vesicle (GSV) translocation to transport GLUT4 to the plasma membrane (6). **B.** Predicted inhibition (blue arrow and nodes) of peroxisome proliferator-activated receptor gamma coactivator 1 alpha (PPARGC1A, PGC-1α, red circle) in sedentary HSA-LR model mice vs WT mice and **C.** Predicted PGC-1α (red circle) activation (orange arrows and nodes) in exercised human skeletal actin long repeat (HSA-LR) model mice vs sedentary HSA-LR model mice. Panel A. created with Biorender.com.

Adenosine monophosphate (AMP)-activated kinase (AMPK) is an intracellular energy sensing protein which is phosphorylated at threonine-172 when adenosine monophosphate (AMP) allosterically binds to it. AMPK can also stimulate AS160 phosphorylation to initiate GLUT4 translocation to the plasma membrane (Arias et al., 2007a). Downstream of AMPK, peroxisome proliferator-activated receptor gamma coactivator 1 alpha (PGC-1α) activates transcription programs for mitochondrial biogenesis, oxidative phosphorylation, and glucose metabolism (Wende et al., 2007a; Cantó and Auwerx, 2009a). Importantly, PGC-1α drives GLUT4 expression and is correlated with improved glucose uptake (Michael et al., 2001; Wende et al., 2007b).

Although exercise confers broad systemic benefits in DM1 models and patients, its effects on insulin sensitivity and GLUT4 translocation in DM1 skeletal muscle remains insufficiently described. Given the important role that AMPK and PGC-1α have in GLUT4 translocation and insulin sensitivity, there is a need for its investigation within the context of DM1 (Cantó and Auwerx, 2009a; b; Michael et al., 2001; Wende et al., 2007b). The human skeletal actin long repeat (HSA-LR) mouse model is a transgenic mouse that models nuclear MBNL aggregation and RNA mis-splicing which translates to myotonia and progressive muscle wasting (Mankodi et al., 2002). Importantly, this model presents with downregulated PGC-1α (Ravel-Chapuis et al., 2018). Although there are no known reports of downregulated PGC-1α in DM1 patients, they do present with dysregulated metabolic dynamics, which could be linked to downregulated PGC-1α (García-Puga et al., 2020, 2022; Mikhail et al., 2022). Thus, we hypothesized that acute activation of the AMPK-PGC-1α axis with an exercise mimetic, 5-aminoimidazole-4-carboxamide ribonucleotide (AICAR), could sensitize the insulin signaling pathway to subsequently improve insulin signaling and GLUT4 translocation in the HSA-LR mouse and a human myotube model of DM1 (Figure 1A).

## 2. Methods

### 2.1 Identification of key molecular pathways

To identify relevant targets within the insulin signaling and exercise pathways in HSA-LR mice, we analyzed proteomic data from tissue of mice that had previously undergone an exercise protocol (Manta et al., 2019). Three to six-month old mice had volitional access to a running wheel for a 7-week period and were sacrificed 24 hours after their last exercise activity. Mass-spectrometry data from quadriceps tissue of 3 sedentary WT mice, 5 sedentary HSA-LR and 5 exercised HSA-LR were entered into Qiagen’s Ingenuity Pathway Analysis (IPA, RRID:SCR_008653, Fall 2025 Release) program to predict protein fold changes and interactions. Raw abundance values were normalized and used to calculate the log_2_ fold change (log_2_FC) to determine the expression change of specific targets. P-values were calculated using the unpaired two-way Welch’s T-test. These values were entered into IPA for subsequent analyses. Using the Core Analysis feature, IPA was used to analyze the proteomic data set against existing public databases and literature to predict the inhibition or activation of upstream pathways. Degree of activation or inhibition of a target or pathway was predicted with a z-score. Z-scores greater than 2 (z>2) indicated activation or upregulation of pathways and targets. Z-scores less than -2 (z<-2) indicated inhibition or downregulation of pathways and targets. A p-value of less than 0.05 (P<0.05) was used to determine statistical significance.

### 2.2 Ethics Approval

Experiments and procedures performed in this study were approved by the University of Ottawa Animal Care Committee and meet the standards of the Canadian Council on Animal Care and Ontario Animals for Research Act. Mice were housed and cared for by the University of Ottawa Animal Care and Veterinary Service.

### 2.3 Animal husbandry and tissue harvesting

Three to six-month old female human skeletal actin-long repeat (HSA-LR) mice (RRID:IMSR_JAX:032031) and wild-type (WT) FVB/N mice (RRID:IMSR_JAX:001800) were subjected to an *ad-libitum* diet and regular light/dark cycles. Mice were injected subcutaneously with 500 mg/kg AICAR (Toronto Research Chemicals, TRC-A611700) or 0.9% saline solution as a control and sacrificed by carbon dioxide administration followed by cervical dislocation 30 minutes after injection. Samples were either snap frozen in liquid nitrogen or mounted in optimal cutting temperature (OCT) mounting media and submerged in liquid nitrogen-cooled isopentane. Samples were stored at -80 °C until use.

### 2.4 Immunofluorescence

OCT mounted tissues were cryosectioned at 10 µm onto slides and stored at -80 °C. Slides were removed from the freezer and allowed to defrost at room temperature for 20 minutes, then fixed in 3.7% formaldehyde for 15 minutes at room temperature. Sections were then washed in 1 x phosphate buffered saline (PBS; Sigma Aldrich, Cat. No. D8537) to remove excess formaldehyde and then circled by a hydrophobic pen before blocking and permeabilization (5% bovine serum albumin (BSA), 5% normal goat serum (NGS), 0.1% Triton-X100) for 30 minutes. Tissue sections were incubated with 1:750 rabbit anti-GLUT4 (Abcam, Cat. No. ab33780, RRID:AB_2191441) and 1:1000 mouse anti-Laminin (Sigma Aldrich, Cat. No. L8271, RRID:AB_477162) in blocking buffer (5% BSA, 5% NGS) overnight at 4 °C in a humidified chamber. The following day, sections were washed 3 x 5 minutes in PBS supplemented with 0.05% tween-20 (PBS-T). Then the sections were incubated in 1:500 goat anti-rabbit AF594 (Thermo Fisher Scientific, Cat. No. A-11012, RRID:AB_2534079) and 1:500 goat anti-mouse AF488 (Thermo Fisher Scientific, Cat. No. RRID:AB_2534088) for 1 hour at room temperature. Afterwards, slides were washed 2 x 5 minutes in PBS-T before adding 0.1 µg/mL Hoechst (Thermo Fisher Scientific, Cat. No. H3570) for 10 minutes at room temperature. Sections were finally washed 2 x 5 minutes in PBS before mounting in Fluoremount-G mounting media (Thermo Fisher Scientific, Cat. No. 00-4958-02). Images were captured on a ZEISS Axio Imager (RRID:SCR_018876) widefield microscope to visualize GLUT4, laminin, and the nucleus in the 594 nm, 488 nm and 350 nm emission wavelengths respectively. Tile images were taken to capture ∼200-300 fibers per sample. Images were analyzed on ZEISS Efficient Navigation (version 3.9, RRID:SCR_013672) imaging software.

### 2.5 Cell culture

Immortalized patient and control myoblast cell lines were maintained at 20% confluence to prevent spontaneous differentiation. Cells were grown at 37 °C in 5% CO2, in Promo Cell Skeletal Muscle Growth Medium (Cat. No. C-23060) which was supplemented with 10% fetal bovine serum, 1% penicillin/streptomycin, and 1% L-glutamine. To passage the cells, media was removed, the cells were washed in 1 x PBS and then incubated in trypsin at 37 °C for 5 minutes before the trypsin was quenched with serum-containing growth media. The cells were then spun down at 1000 RPM for 5 minutes, resuspended in growth media and placed into a T75 flask. If cells were directed toward experimentation, they were counted and seeded into a collagen (Gibco Collagen I rat tail, A1048301) coated dish to ensure cell adherence during differentiation. Cells were grown to 90-100% confluence then switched to differentiation media (DM) (Dulbecco’s Modified Eagle Medium (DMEM), 2% horse serum, 1% penicillin/streptomycin, and 1% L-glutamine). The cells were maintained in DM for 4 days before drug treatments.

### 2.6 Cell drug treatments

We used and experimented on DM1 patient-derived myotubes that were obtained as a gift from Dr. Elena Pegoraro’s group (Pantic et al., 2016). Information pertaining to the sex, age, and biopsy location is described in Table 1.

**Table 1.** Cell model information. Patient’s sex, age, and biopsy site.

| Cell Line | Sex | Age | Biopsy Site |
| --- | --- | --- | --- |
| Control-1 | Male | 41 | *Vastus Lateralis* |
| Control-3 | Male | 36 | *Vastus Lateralis* |
| DM1-1 | Female | 29 | *Biceps Brachii* |
| DM1-2 | Male | 19 | *Vastus Lateralis* |

Myoblasts were plated on 6 well plates and differentiated for 4 days before serum starvation for 4 hours in DMEM. Myotubes were then treated with 2mM AICAR in DMSO for 1 hour followed by 100 nM insulin in HEPES buffer for 30 minutes at 37 °C. Vehicle treatment (Veh) cells were treated with a combination of 2% DMSO control and 1.25 µM HEPES buffer. Cells were treated with either vehicle, AICAR, insulin, or both AICAR and insulin.

### 2.7 Western blot

Skeletal muscle tissues were lysed in RIPA buffer (150 mM NaCl, 50 mM Tris HCl, 1% Triton-X100, 0.1% SDS, 0.5% sodium deoxycholate) supplemented with protease and phosphatase inhibitor (Sigma Aldrich, Cat. No. PPC1010-1ML). The tissue was lysed with 3.2 mm steel beads and placed into the Qiagen Tissue Lyser at 25 Hz for 3 x 90 seconds, with 2 minutes cooling on ice in between cycles. Lysates were then placed on a rotator at 4 °C for 2 hours before being spun down at 12 000 RPM for 20 minutes. Supernatant was collected and lysates were stored at -80 °C. Protein was quantified using a DC assay kit (Bio-Rad, Cat. No. 5000112) alongside a BSA standard that ranged from 0.125 to 2 µg/mL.

Myotubes were lysed in the same RIPA buffer cocktail on ice and scraped off the 6-well plate with a cell scraper. Lysates were vortexed and then placed on a rotator at 4 °C for 30 minutes before being spun down at 12 000 RPM for 20 minutes. Supernatant was collected and lysates were stored and quantified the same way that muscle lysates were.

Samples were prepared with 4X Laemmli buffer (Bio-Rad, Cat. No. 161-0747) and supplemented with 2-mercaptoethanol. Samples were then placed on a heat block at 95 °C for 5 minutes before being run through a 7.5% polyacrylamide stacking and 8% polyacrylamide resolving gel in tris-glycine running buffer (25 mM Tris-Base, 190 mM glycine, 0.1% SDS). The samples were run for 15 minutes at 115 V and then 75 minutes at 125 V. The samples were then transferred to 0.2 µm PVDF (Bio-Rad, Cat. No. 1620177) in a cold tris-glycine transfer buffer (25 mM Tris-Base, 190 mM glycine, 20% ethanol) for 90 minutes at 100 V at 4 °C. Membranes were then blocked in Licor blocking buffer (Licor, Cat. No. 927-60001) for 1 hour at room temperature. Primary antibodies (in Licor blocking buffer, 0.1% Tween), detailed in Supplemental Table 1, were added overnight at 4 °C on a rocker (Supplemental Table 1). The following day blots were washed in tris buffered saline – tween (TBS-T; 20 mM Tris-HCL, 150 mM NaCl, 0.1% Tween-20) for 3 x 5 minutes before being incubated in the respective secondary antibody for 1 hour at room temperature on a rocker (Supplemental Table 1). Finally, blots were incubated in Clarity Max enhance chemiluminescent substrate (Bio-Rad, Cat. No. 1705062) and bands imaged on the ChemiDoc Imaging System (Bio-Rad, RRID:SCR_019037). Band densitometry was analyzed on Image Lab software (Bio-Rad, RRID:SCR_014210) and all proteins were normalized to vinculin loading protein.

### 2.8 Statistical analyses

Statistical comparisons were made once the data sets were assessed for normality with the Shapiro-Wilk test. If datasets failed (p-value<0.05), then they were analyzed with the unpaired one-way Welch’s t-test. Otherwise, the tests were conducted with unpaired one-way Student’s t-test. Multiple comparisons were analyzed with two-way ANOVA followed by Tukey’s post hoc analysis. Statistical significance was determined with a threshold of p< 0.05. Grubb’s extreme student deviate test was used to determine and exclude outliers. Statistics were performed, and graphs were generated using Prism GraphPad (RRID:SCR_002798) with graphs depicted as mean +/-standard error of the mean (SEM).

## 3. Results

### 3.1 Ingenuity Pathway Analysis tool predicts PGC-1α and AMPK to be dysregulated in HSA-LR mouse model

There were 882 targets identified in the sedentary HSA-LR mice versus sedentary WT condition and 884 targets identified in the exercised HSA-LR mouse versus sedentary HSA-LR mouse condition (Supplemental Table 2,3). Of the targets, solute carrier family 2 member 4 (SLC2A4), also known as GLUT4, was upregulated in the exercised HSA-LR condition (Table 2, Supplemental Table 2,3).

**Table 2.** SLC2A4 expression measures in sedentary and exercised HSA-LR conditions.

| Condition | Expression Log Ratio | Expression p-value |
| --- | --- | --- |
| Sedentary HSA-LR vs sedentary WT | 2.82 | 6.0x10<sup>-2</sup> |
| Exercised HSA-LR vs sedentary HSA-LR | 3.80 | 4.0x10<sup>-2</sup> |

When assessing the predicted Canonical pathways, PI3K-Akt was predicted to be activated in sedentary and exercised conditions (Table 3, Supplemental Table 4, 5). Within the pathway, the predicted activated molecules were not PI3K or Akt, but rather targets downstream of these molecules (Supplemental Table 6,7). Of these downstream effectors overlapping with the proteomic dataset, none of them met the thresholds for z-score (>|2|) and p-value (<0.05) (Supplemental Table 6,7).

**Table 3.** PI3K-Akt canonical pathway activation measures in sedentary and exercised HSA-LR conditions.

| Condition | Activation z-score | P-value |
| --- | --- | --- |
| Sedentary HSA-LR vs sedentary WT | 2.67 | $1.05 \times 10^{-4}$ |
| Exercised HSA-LR vs sedentary HSA-LR | 2.18 | $1.08 \times 10^{-4}$ |

Similarly, the GLUT4 translocation pathway was increased in both conditions. However, within the GLUT4 translocation pathway, GLUT4 was significantly upregulated only in the exercised HSA-LR condition (Table 2, 4, Supplemental Table 4,5,8,9).

For the predicted upstream regulators, INSR (insulin receptor) and PPARGC1A (PGC-1α) appeared as relevant targets as both are implicated in as glucose metabolism regulators. INSR was predicted to be inhibited in the sedentary HSA-LR mice, but was activated in the exercised HSA-LR mice (Table 5, Supplemental table 10, 11). Similarly, PGC-1α was predicted to be inhibited in sedentary HSA-LR mice and was upregulated in exercised HSA-LR mice (Figure 1B – C, Table 6, Supplemental Table 10, 11).

**Table 4.** Translocation of SLC2A4 (GLUT4) canonical pathway activation measures in sedentary and exercised HSA-LR conditions.

| Condition | Activation z-score | P-value |
| --- | --- | --- |
| Sedentary HSA-LR vs sedentary WT | 2.84 | $9.05 \times 10^{-9}$ |
| Exercised HSA-LR vs sedentary HSA-LR | 2.84 | $9.33 \times 10^{-9}$ |

**Table 5.** INSR (insulin receptor) predicted activation measures in sedentary and exercised HSA-LR conditions.

| Condition | Activation z-score | P-value |
| --- | --- | --- |
| Sedentary HSA-LR vs sedentary WT | -4.27 | $2.67 \times 10^{-60}$ |
| Exercised HSA-LR vs sedentary HSA-LR | 5.35 | $3.56 \times 10^{-60}$ |

**Table 6.** PPARGC1A (PGC-1α) upstream regulator predicted activation measures in sedentary and exercised HSA-LR conditions.

| Condition | Activation z-score | P-value |
| --- | --- | --- |
| Sedentary HSA-LR vs sedentary WT | -2.59 | $3.75 \times 10^{-48}$ |
| Exercised HSA-LR vs sedentary HSA-LR | 6.51 | $4.61 \times 10^{-48}$ |

### 3.2 AMPK-PGC-1α axis has impaired response to AICAR in HSA-LR mice

To determine the specific effect of AMPK stimulation on PGC-1α expression and insulin signaling molecules, mice were subjected to an acute AICAR treatment. They received a subcutaneous injection with 500 mg/kg AICAR or saline. The mice were sacrificed, and tissues were collected 30 minutes after injection to capture the transient action of AMPK. Samples were analyzed with western blotting to assess the abundance and activity levels of proteins involved in the key signaling pathways (Figure 2A, Supplemental Figure 3-6, 9). At baseline, HSA-LR mice expressed lower total AMPK, but not phosphorylated threonine-172 AMPK (pAMPK) compared to WT mice. No difference was observed when pAMPK was normalized to total AMPK (Figure 2B-D). Analysis indicated no significant difference in baseline expression of PGC-1α between HSA-LR and WT groups (Figure 2E). However, there was significantly less GLUT4 in the HSA-LR group compared to the WT group (Figure 2F).

**Figure 2.**
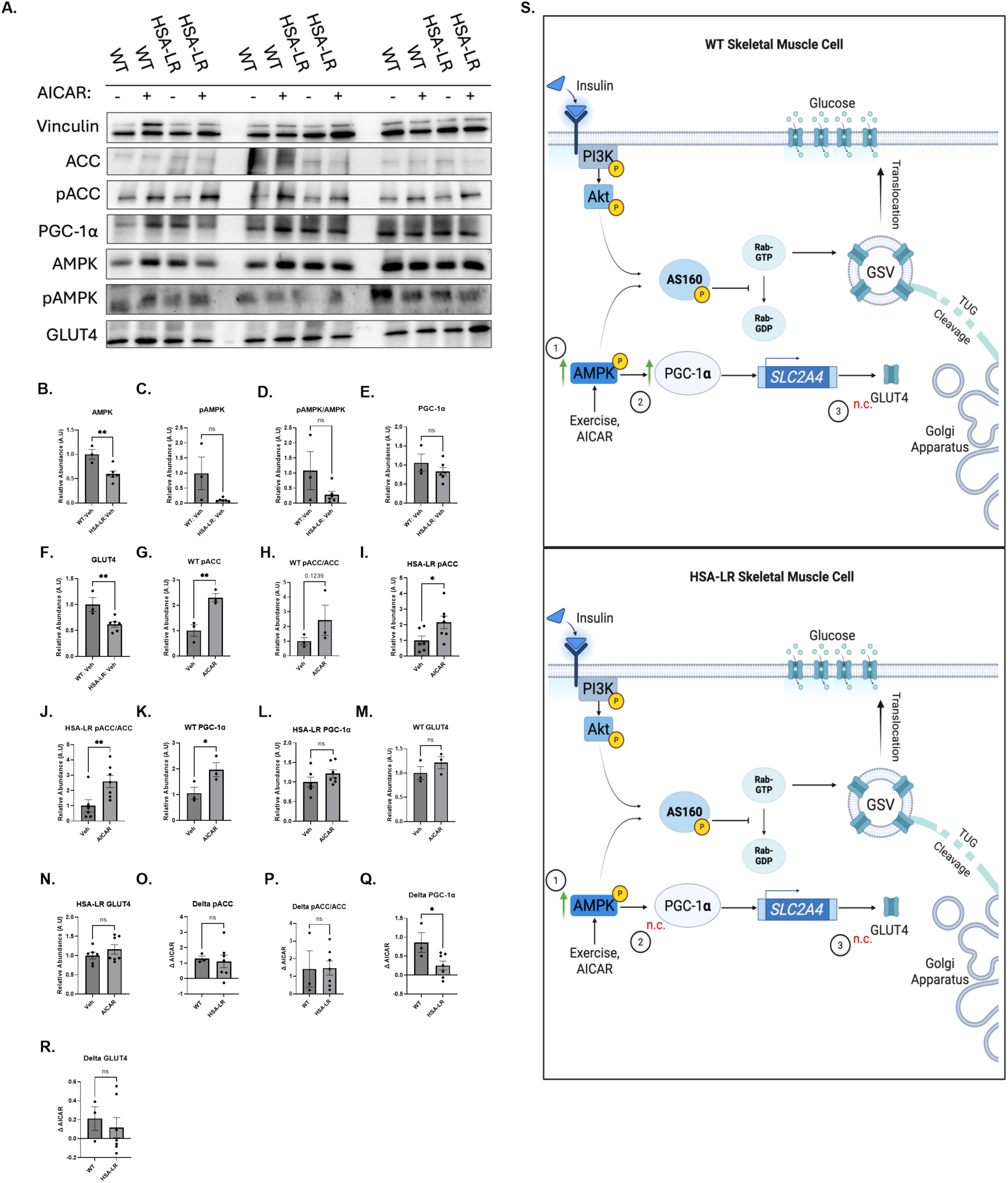
AMPK-PGC-1α pathway in WT and HSA-LR mice. **A.** Representative western blots (whole blots found in Supplemental Figure 3-6, 9) of proteins along the exercise signaling pathway in gastrocnemius tissue of wild-type (WT) and human skeletal actin long repeat (HSA-LR) mice injected with saline (-) or 5-aminoimidazole-4-carboxamide ribonucleotide (AICAR, +). **B.** Decreased levels of adenosine monophosphate activated protein kinase (AMPK), **C.** but not threonine-172 phosphorylated AMPK (pAMPK) in HSA-LR mice. No difference when **D.** pAMPK is normalized to total AMPK (pAMPK/AMPK). **E.** Comparison of peroxisome proliferator-activated receptor gamma coactivator 1 alpha (PGC-1α) protein levels shows no significant difference between WT and HSA-LR mice at baseline **F.** Comparison of glucose transporter 4 (GLUT4) protein levels shows higher protein in WT compared to HSA-LR mice at baseline. **G. & H.** Increase of the phosphorylated serine-79 acetyl-CoA carboxylase (pACC) and pACC/ACC in WT mice after AICAR treatment. **I. & J.** Increase of the pACC and pACC/ACC in HSA-LR mice after AICAR treatment. **K.** Increase of the PGC-1α in WT mice after AICAR treatment. **L.** Increase of PGC-1α from HSA-LR mice after AICAR treatment. **M.** No change in GLUT4 levels in WT mice after AICAR treatment. **N.** No change in GLUT4 levels in HSA-LR mice after AICAR treatment **O. & P.** Delta values (treatment values – baseline values) with respect to AICAR treatment for pACC, pACC/ACC shows no differences in the responses to AICAR for WT and HSA-LR mice. **Q.** Delta values with respect to AICAR treatment for PGC-1α indicate a greater response in WT compared to HSA-LR. **R.** Delta values with respect to AICAR treatment for GLUT4 shows no differences between WT and HSA-LR mice**. S.** Graphical depiction of the molecular response to AICAR treatment in WT and HSA-LR mice. Green arrows indicate an increased level of the target after AICAR treatment. Increased signaling of AMPK (1) as reflected in the increased pACC levels (not shown), increased PGC-1α (2) in WT mice, but no change (red n.c.) in HSA-LR, and GLUT4 (3) reflects no change (n.c.) in either WT or HSA-LR mice. Panel S. created by Biorender.com. N = 3 WT (Veh), 3 WT (AICAR), 6 HSA-LR (Veh), 7 HSA-LR (AICAR) mice. All samples run in technical duplicates. One-tailed unpaired Welch’s T-test for panels C, M, N, O, P, R. Remaining panels analyzed with one-tailed unpaired Student’s T-test. *p<0.05, **p < 0.005, ***p<0.0005. Graphs are means +/-SEM.

After AICAR stimulation, AMPK activity showed no changes in WT or HSA-LR mice (Supplemental Figure 1A -F). To validate the effect of AICAR, acetyl-CoA carboxylase (ACC) was analyzed for phosphorylation on serine-79, a stable readout for AMPK activity (Galic et al., 2018). There was no change in total ACC in either WT or HSA-LR mice (Supplemental Figure 1G - I). However, there was a marked increase in the pACC levels after AICAR treatment, reflecting adequate stimulation of the AMPK signaling pathway (Figure 2G-J).

PGC-1α levels increased in WT mice after AICAR treatment (Figure 2K), but not in HSA-LR mice (Figure 2L). There were also no significant differences in GLUT4 expression after treatment in WT or HSA-LR animals (Figure 2M, N). We then calculated the delta AICAR values for each target for the experimental groups (treatment group values – baseline group values) to compare the normalized response to AICAR treatment. This comparison elucidates the proportional response of the pathway targets. Delta AICAR comparisons for pACC yielded no differences, further validating the proportional activation of AMPK signaling with AICAR in both experimental groups (Figure 2O, P). Delta AICAR comparisons of PGC-1α showed that HSA-LR mice had an impaired response to AICAR treatment (Figure 2Q). Further, delta AICAR comparisons for GLUT4 showed no differences in responses to treatment (Figure 2R). In summary, we observed that even with proportional AICAR activation of AMPK, we saw that PGC-1α response was impaired in HSA-LR mice (Figure 2S).

### 3.3 Akt-AS160 has impaired response to AICAR in HSA-LR mice

Next, we wanted to determine the impact AICAR had on intrinsic insulin signaling. Thus, we assessed phosphorylated serine-473 Akt (pAkt) and phosphorylated threonine-642 AS160 (pAS160) on western blot (Figure 3A, Supplemental Figure 7-9). No significant differences were detected in baseline pAkt, pAkt/Akt or pAS160 between WT and HSA-LR mice (Figure 3B - D). After AICAR treatment, pAkt and pAkt/Akt showed an increased response in the WT mice, but this was not reflected in the HSA-LR mice (Figure 3E-H). When total Akt was assessed, there were no differences in total expression before or after AICAR treatment (Supplemental Figure 1J -L). Downstream of Akt, pAS160 was measured to determine the signaling capacity for GLUT4 translocation. In WT animals, AICAR stimulation led to a significant increase in pAS160 (Figure 3I). However, there was no such trend found in the HSA-LR mice (Figure 3J).

**Figure 3.**
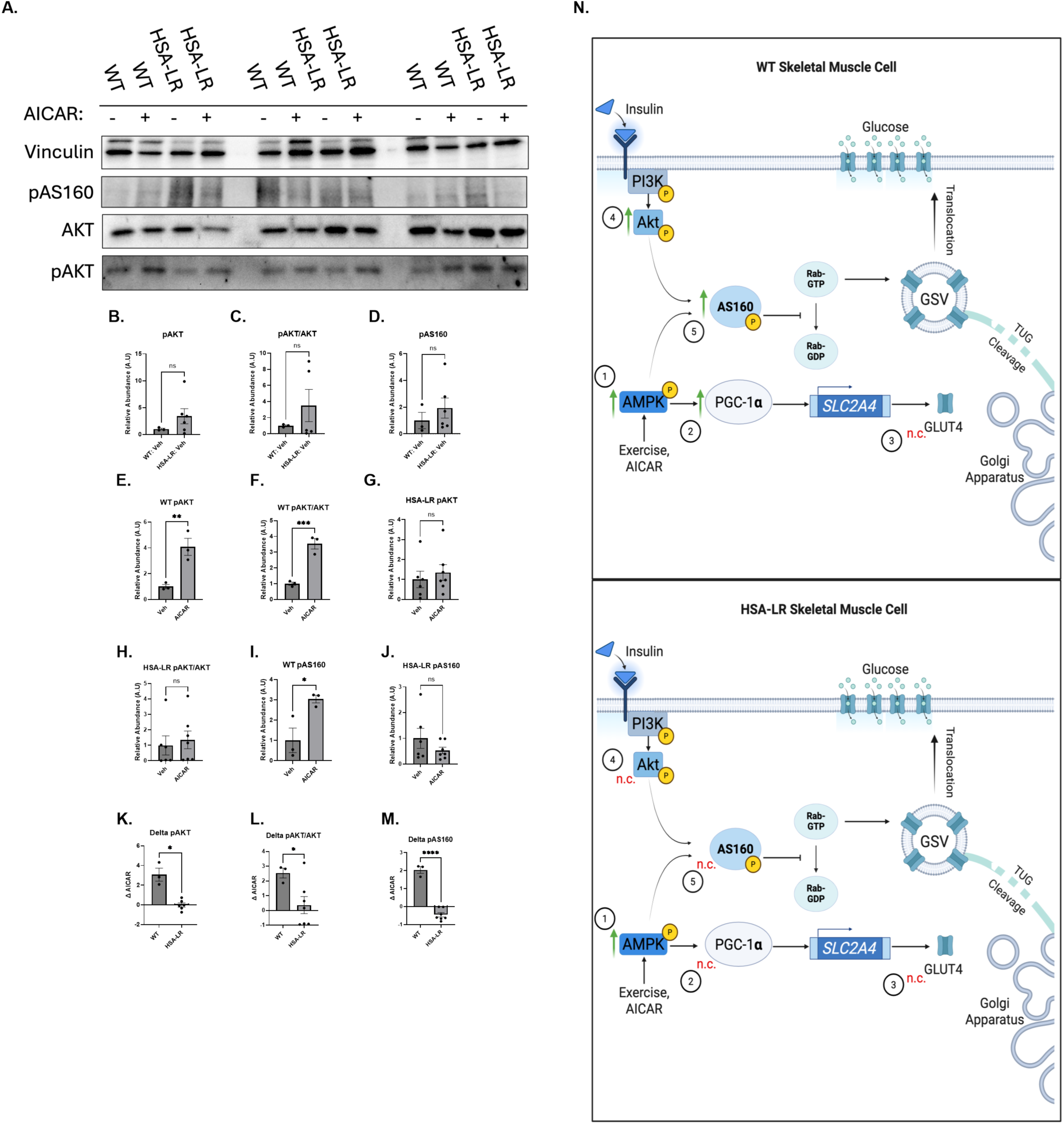
Akt-AS160 pathway in WT and HSA-LR mice. **A.** Representative western blots (whole blots found in Supplemental Figure 7-9) of proteins along the insulin signaling molecular pathways in gastrocnemius tissue of wild-type (WT) and human skeletal actin long repeat (HSA-LR) mice injected with saline (-) or 5-aminoimidazole-4-carboxamide ribonucleotide (AICAR, +). **B. & C.** Comparison of pAkt and pAkt/Akt protein levels shows no significant difference between WT and HSA-LR mice. **D.** Comparison of phosphorylated threonine-642 Akt substrate of 160 kDA (pAS160) protein levels shows higher total protein in WT compared to HSA-LR mice. **E. & F.** Increase of the pAkt and pAkt/Akt after AICAR treatment in WT mice. **G. & H.** Quantification of the pAkt and pAkt/Akt observed in the HSA-LR mice. **I.** Increase of pAS160 observed in the WT mice after AICAR injection. **J.** Quantification of the pAS160 in HSA-LR mice. **K.-M.** Quantification of the delta values with respect to AICAR treatment for pAkt, pAkt/Akt, pAS160. **N.** Graphical depiction of the response of molecules after treatment with AICAR in WT and HSA-LR mice. Green arrows indicate an increased level of the target after AICAR treatment. (4) indicates the increased phosphorylation of Akt in WT mice with no change (n.c.) observed in HSA-LR mice, reflecting the increased pAS160 (5) in WT mice, but no change (n.c.) in HSA-LR. N = 3 WT (Veh), 3 WT (AICAR), 6 HSA-LR (Veh), 7 HSA-LR (AICAR) mice. Panel N. created by Biorender.com. All samples run in technical duplicates. All samples run in technical duplicates. One-tailed unpaired Welch’s T-test for panels B, C, G, H, K, L. Remaining panels analyzed with one-tailed unpaired Student’s T-test. *p<0.05, **p < 0.005, ***p<0.0005. Graphs are means +/-SEM.

Again, delta AICAR values were calculated to compare the proportional changes of protein expression after AICAR treatment. The delta AICAR for pAkt revealed that HSA-LR mice had a significantly impaired response to AICAR compared to the WT mice (Figure 3K, L). Similarly, delta AICAR for pAS160 revealed that pAS160 change was significantly lower in HSA-LR mice compared to WT mice (Figure 3M). Despite proportional AMPK stimulation in WT and HSA-LR mice, the insulin signaling pathway was not adequately stimulated in the HSA-LR mice (Figure 3N).

### 3.4 Immunofluorescence indicates trends of increased GLUT4 translocation in WT mice after AICAR stimulation

In gastrocnemius tissue from vehicle- and AICAR-injected WT and HSA-LR mice, GLUT4 translocation was assessed by probing for GLUT4 and laminin. Localization was quantified with the proportion of the total GLUT4 and GLUT4 colocalizing with the laminin-labeled plasma membrane (Figure 4A). The pattern of the GLUT4 staining in the vehicle-treated WT mice was dispersed in the cytosol. Some of the GLUT4 staining was found along the plasma membrane region, denoted by the laminin staining. HSA-LR mice exhibited higher membrane GLUT4 staining after AICAR treatment, but these differences were lost when normalized to total GLUT4 staining (Supplemental Figure 1M – P). WT mice treated with AICAR had trends (p = 0.0573) indicating increased GLUT4-Laminin:Total GLUT4, suggesting increased levels of GLUT4 at the plasma membrane (Figure 4B). HSA-LR mice did not show (p = 0.26) any shifts in the GLUT4-Laminin:Total GLUT4 after AICAR treatment (Figure 4C). When the delta AICAR was compared there were no differences found, indicating no significant differences in relative GLUT4 translocation (Figure 4D).

**Figure 4.**
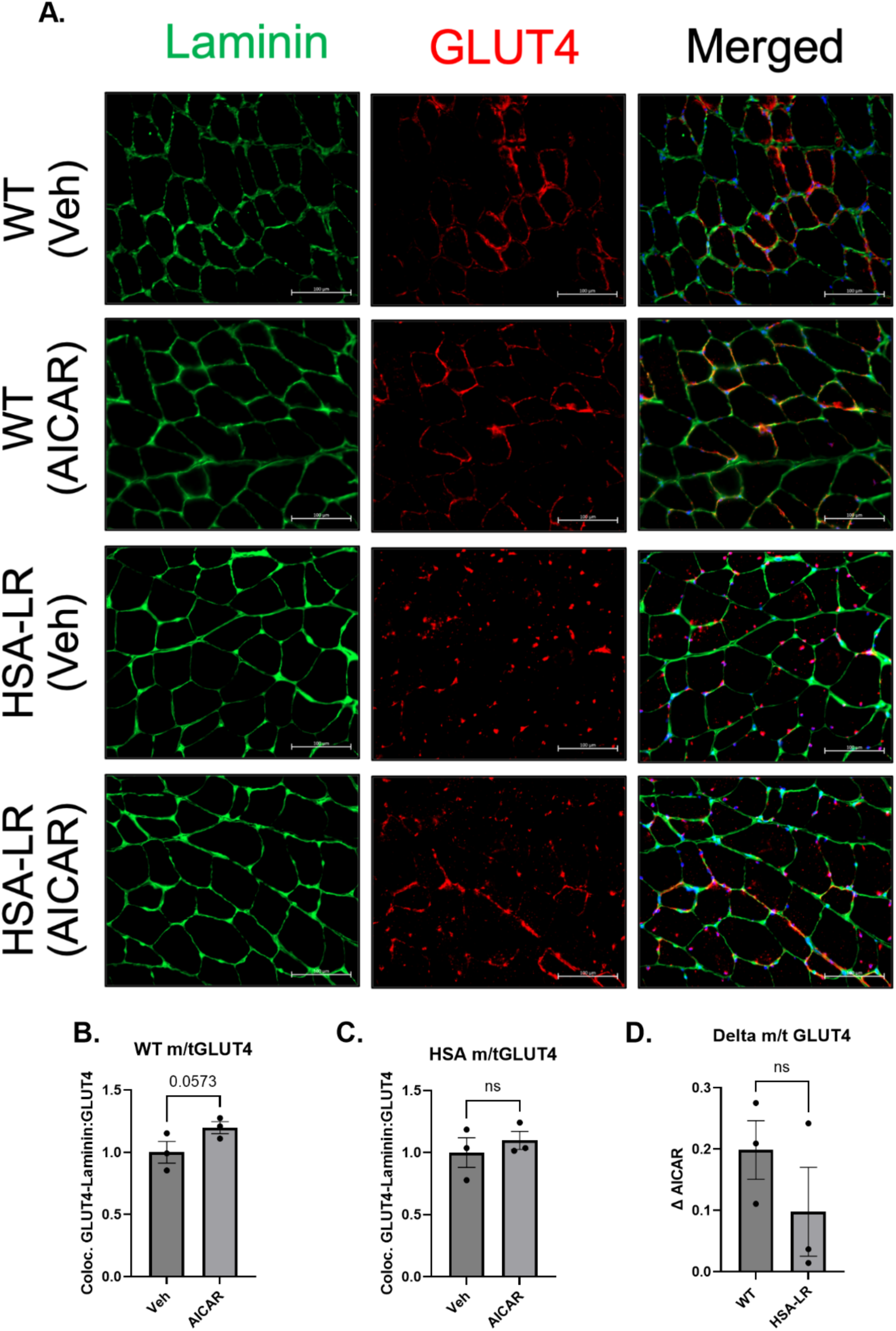
Immunofluorescence of GLUT4 labelling. **A**. Representative images of immunofluorescent staining of laminin (green), glucose transporter 4 (GLUT4, red) and 4′,6-Diamidino-2-phenylindole (DAPI, blue, merged image) in gastrocnemius sections from saline vehicle (Veh) treated and 5-Aminoimidazole-4-carboxamide ribonucleoside (AICAR) treated mice. **B.** A quantification of the GLUT4-Laminin colocalization (mGLUT4) pixel area relative to the total GLUT4 (tGLUT4) pixel area in WT sections. **C**. A quantification of the GLUT4-Laminin colocalization pixel area relative to the total GLUT4 (tGLUT4) pixel area in HSA-LR sections. **D.** A comparison of the delta responses to AICAR treatment in WT compared to HSA-LR mice. N = 3 WT (Veh), 3 HSA-LR (Veh), 3 WT (AICAR), 3 HSA-LR (AICAR). Experiment conducted in technical duplicates and 200-300 fibres analyzed per section. Scale bar is 100 μm. One-tailed unpaired Student’s T-test *p<0.05, **p < 0.005, ***p<0.0005. Graphs are means +/-SEM.

### 3.5 AMPK axis is stimulated in human control and DM1 myotubes

Healthy control (CTL) cells and DM1 myotubes were treated with insulin alone, AICAR alone, or AICAR and insulin combined, and then probed for targets to assess the impact that treatment had on the AMPK signaling pathway (Figure 5A, Supplemental Figure 10-13, 16). At baseline, total AMPK was reduced in DM1 myotubes (Figure 5B). However, phosphorylated threonine-172 AMPK (pAMPK) in DM1 myotubes was greater in DM1 myotubes (Figure 5C,D). There was no change in pAMPK after AICAR or insulin treatments (Figure 5E). Given the transient nature of AMPK, ACC, a stable downstream target of AMPK, was also assessed. Phosphorylated serine-79 ACC (pACC) increased with AICAR and combination treatments in CTL and DM1 myotubes (Figure 5F). There were no differences in PGC-1α or GLUT4 expression prior to or after treatments (Figure 5G – H).

**Figure 5.**
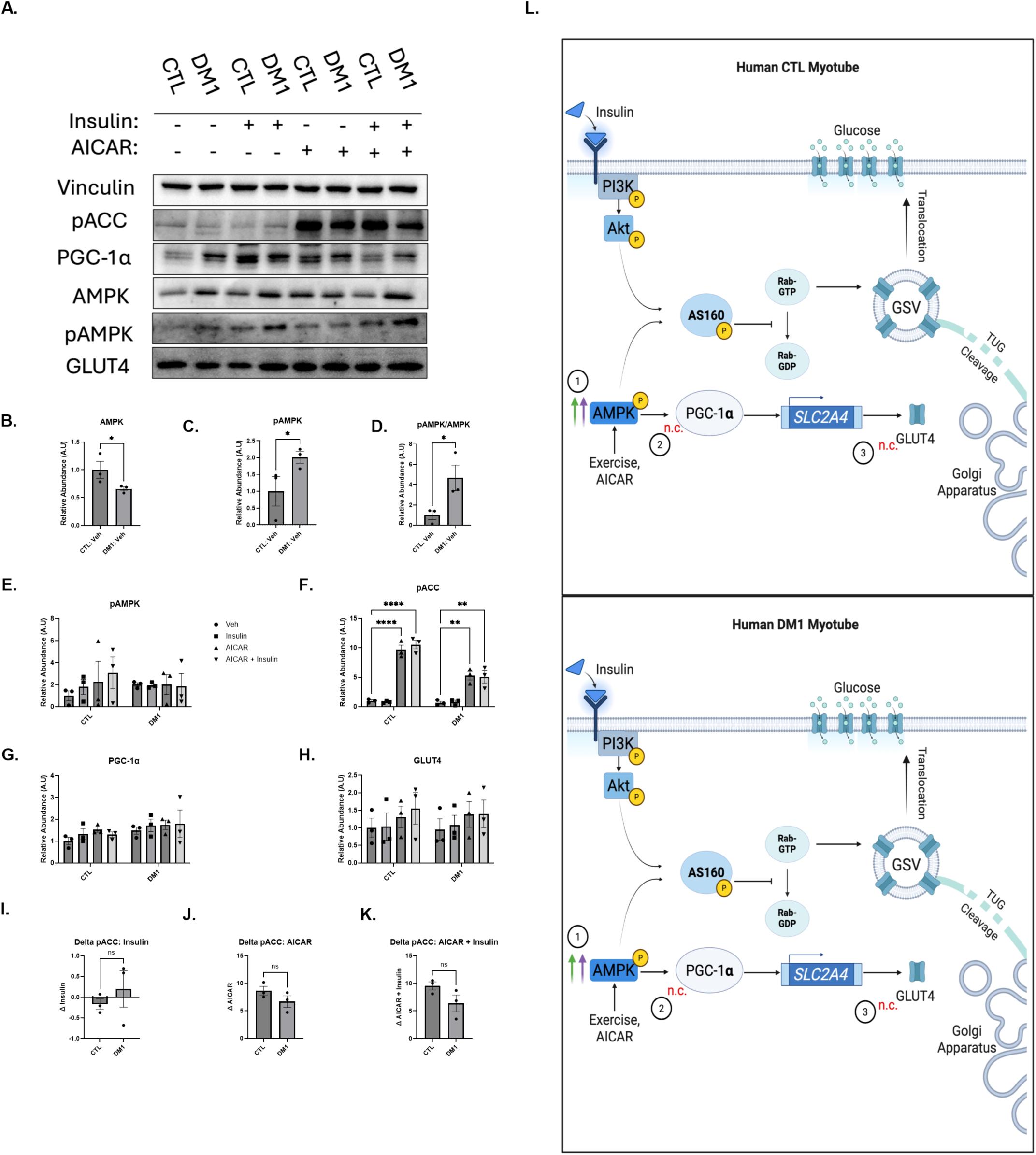
AMPK-PGC-1α pathway in CTL and DM1 myotubes. **A.** Representative western blots (whole blots found in Supplemental Figure 10-13, 16) of targets along the exercise molecular pathways in human control (CTL) and myotonic dystrophy type 1 (DM1) myotubes mice treated with vehicle (-), insulin, 5-aminoimidazole-4-carboxamide ribonucleotide (AICAR), or a combination of insulin and AICAR. **B.** At baseline, total adenosine monophosphate-activated protein kinase (AMPK) is decreased in human DM1 myotubes. **C.** At baseline, phosphorylated threonine-172 AMPK (pAMPK) is increased in human DM1 myotubes. **D.** At baseline, normalized pAMPK (pAMPK/AMPK) is increased in human DM1 myotubes. **E.** No change for the pAMPK observed in the human CTL and DM1 myotubes after insulin, AICAR or combination treatment. **F.** Increase of the phosphorylated serine-79 acetyl-CoA carboxylase (pACC) observed in the human CTL and DM1 myotubes after AICAR treatment. **G.** No change for the PGC-1α observed in the human CTL and DM1 myotubes after insulin, AICAR or combination treatment. **H.** No change for the glucose transporter 4 (GLUT4) observed in the human CTL and DM1 myotubes after insulin, AICAR or combination treatment. **I.-K.** No change of the delta values with respect to **I.** insulin, **J.** AICAR, or **K.** insulin + AICAR treatment for pACC between CTL and DM1 cell lines. **L.** Graphical depiction of pathway proteins responding to AICAR (green arrow) and/or insulin + AICAR combination (purple arrow) in CTL and DM1 myotubes. (1) Indicates the increased signaling of AMPK levels, (2) reflects no change (n.c.) of PGC-1α and (3) glucose transporter 4 (GLUT4) in DM1 or CTL myotubes. Panel L. created by Biorender.com. N = 2 CTL, 2 DM1. One way ANOVA followed by Tukeys post-hoc test for multiple comparisons E - H., One-tailed unpaired Student’s T-test *p<0.05, **p < 0.005, ***p<0.0005. Graphs are means +/-SEM.

Like the analysis in the mice, delta AICAR, delta insulin and delta combination values were calculated to compare the proportional changes of protein expression after treatments. For pACC, there were no significant differences found after insulin, AICAR or combination treatments (Figure 5I-K), further validating the proportional activation of AMPK. In summary, despite AMPK stimulation there were no observed changes in PGC-1α or GLUT4 expression (Figure 5L).

### 3.6 Akt-AS160 signaling is impaired in DM1 myotubes

CTL and DM1 myotubes were treated with vehicle, insulin, AICAR or a combination of both and then analyzed for Akt and AS160 signaling (Figure 6A, Supplemental Figure 2G – J, 14-16). DM1 myotubes had significantly higher pAkt expression than CTL at baseline (Figure 6B). Unlike CTL myotubes, DM1 pAkt showed no significant response to insulin treatment, but did exhibit an increase with combination treatment (Figure 6C). Similar trends were observed after normalization to total Akt (Figure 6D), with a loss of statistical significance for combination treatment (Figure 6E).

**Figure 6.**
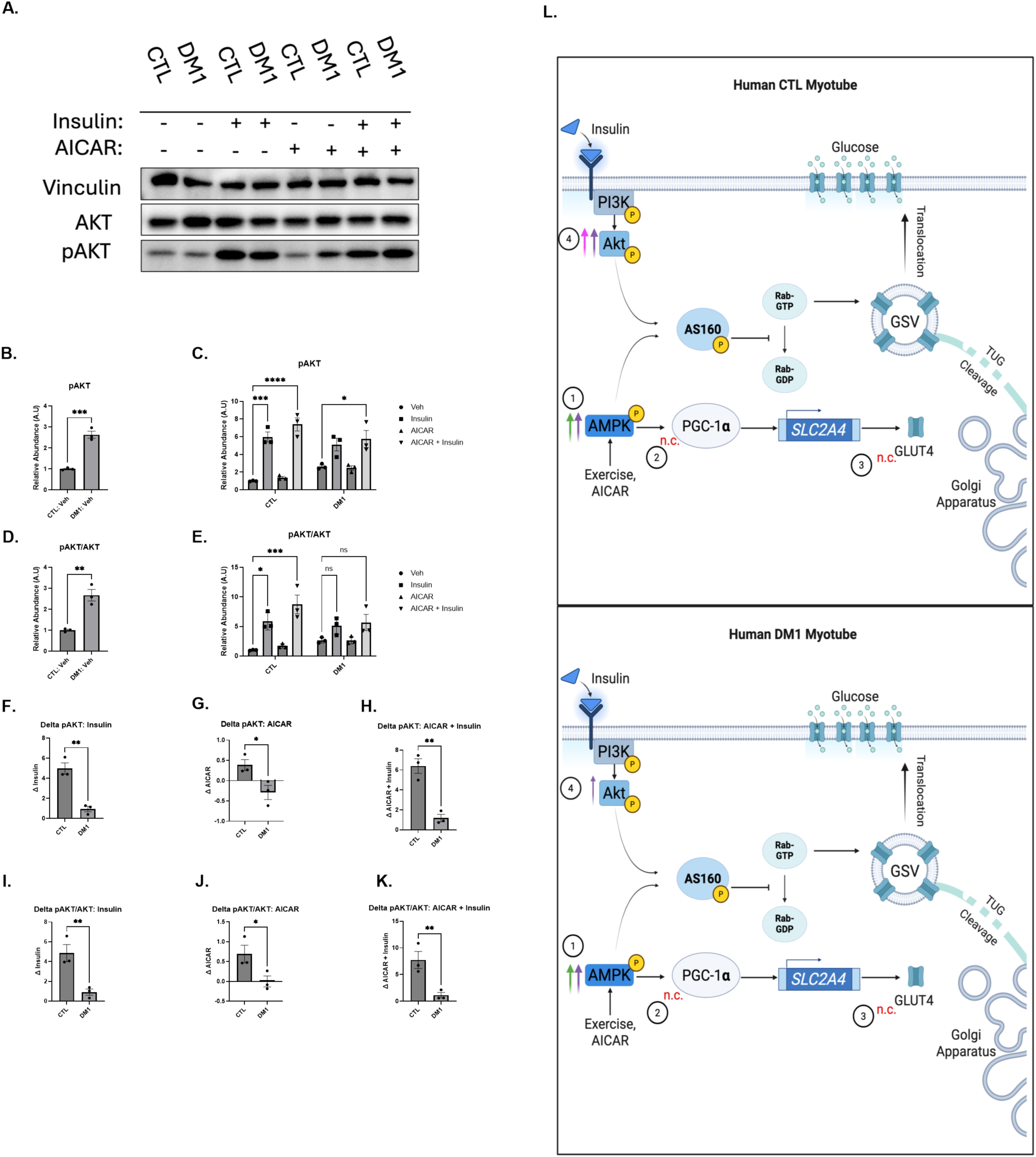
pAkt/Akt signaling in CTL and DM1 myotubes. **A.** Representative western blots (whole blots found in Supplemental 14,16) of Akt and pAkt in human control (CTL) and myotonic dystrophy type 1 (DM1) myotubes treated with vehicle (-), insulin, 5-Aminoimidazole-4-carboxamide ribonucleoside (AICAR), or a combination of insulin + AICAR. **B.** Increase of the pAkt observed in the human CTL after insulin alone and combination treatment and with an increase only after combination treatment seen in DM1 myotubes. **C.** Increased levels of pAkt in vehicle treated DM1 (DM1:Veh) compared to vehicle treated control (CTL:Veh) myotubes. **D.** Quantification of the pAkt/Akt observed in the human CTL and DM1 myotubes. **E.** Comparison of pAkt/Akt levels shows increased levels in vehicle treated DM1 (DM1:Veh) compared to vehicle treated control (CTL:Veh) myotubes. **F.-H.** Quantification of the delta values with respect to **F.** insulin, **G.** AICAR, or **H.** insulin + AICAR treatment for pAkt reflecting greater delta values in the CTL myotubes. **I.-K.** Greater delta values with respect to **I.** insulin, **J.** AICAR, or **K.** insulin + AICAR treatment for pAkt/Akt in CTL myotubes. **L.** Graphical depiction of the response of molecules after treatment with insulin (pink arrow), AICAR (green arrow) or insulin + AICAR (purple arrow). (4) Reflects increased phosphorylated Akt ratio in CTL myotubes and relatively smaller increase in DM1 myotubes. Panel L. created by Biorender.com. N = 2 CTL, 2 DM1. One way ANOVA followed by Tukeys post-hoc test for multiple comparisons panels C. & E. One-tailed Welch’s T-test for panel J. One-tailed unpaired Student’s T-test *p<0.05, **p < 0.005, ***p<0.0005. Graphs are means +/-SEM.

Again, delta insulin, delta AICAR and delta combination values were calculated to compare the proportional changes of protein expression after treatments. DM1 myotubes exhibited an impaired pAkt and pAkt/Akt response to insulin, AICAR, and combination treatments (Figure 6F-K). Altogether, there is consistent indication of impaired Akt signaling in DM1 myotubes (Figure 6L).

AS160 phosphorylation levels were also measured in CTL and DM1 myotubes (Figure 7A). Consistent with pAkt and pAkt/Akt, we found that baseline pAS160 was higher in DM1 myotubes compared to CTL myotubes (Figure 7B). Both CTL and DM1 myotubes exhibit pAS160 increase after combination treatment (Figure 7C). When pAS160 was normalized to total AS160 (pAS160/AS160), similar trends were observed (Figure 7D -E).

**Figure 7.**
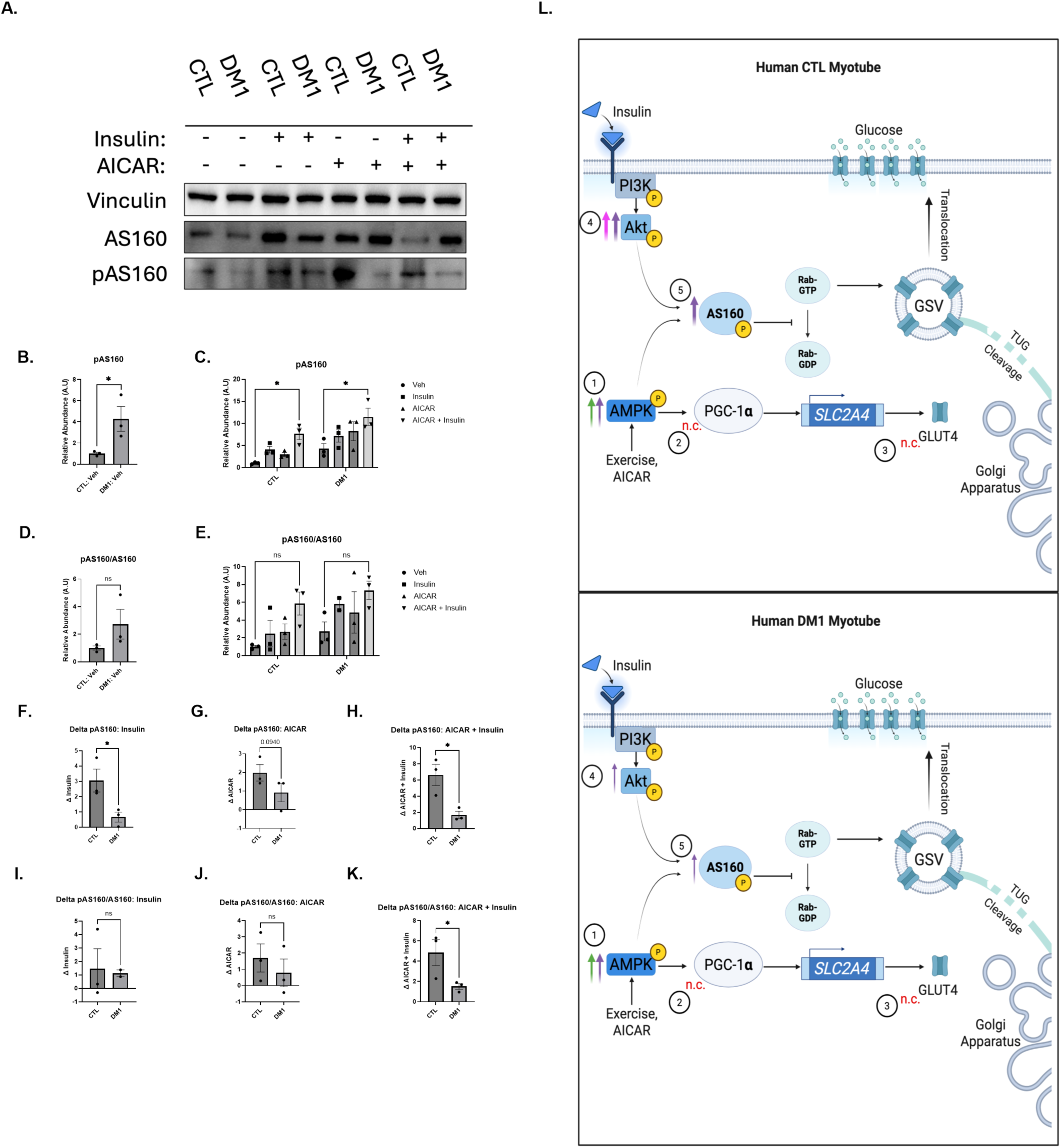
AS160 signaling in CTL and DM1 myotubes. **A.** Representative western blots (whole blots found in Supplemental 15,16) of the pAS160 and AS160 in human CTL and DM1 myotubes treated with vehicle (-), insulin, 5-aminoimidazole-4-carboxamide ribonucleotide (AICAR), or a combination of insulin and AICAR. **B.** Quantification of the pAS160 observed in the human CTL and DM1 myotubes. **C.** Comparison of pAS160 levels shows increased levels in vehicle treated DM1 (DM1:Veh) compared to vehicle treated control (CTL:Veh) myotubes. **D.** No significant change of the pAS160/AS160 observed in the human CTL and DM1 myotubes after treatments. **E.** Comparison of pAS160/AS160 levels shows no significant difference between vehicle treated DM1 (DM1:Veh) compared to vehicle treated control (CTL:Veh) myotubes. **F.-H.** Greater CLT delta values with respect to **F.** insulin, **G.** AICAR, or **H.** insulin + AICAR treatment for pAS160. **I.-K.** Quantification of the delta values with respect to **I.** insulin, **J.** AICAR, or **K.** insulin + AICAR treatment for pAS160/AS160 with a greater delta value observed in the CTL myotubes. **L.** Graphical depiction of the response of molecules after treatment with insulin (pink arrow), AICAR (green arrow) or insulin + AICAR (purple arrow). (5) reflects increased phosphorylation of AS160 (dashed arrow) in CTL and DM1 myotubes after AICAR and insulin treatment with no change (n.c.) seen in DM1 myotubes. Panel L. created by Biorender.com. N = 2 CTL, 2 DM1. One way ANOVA followed by Tukeys post-hoc test for multiple comparisons C. & E. One-tailed unpaired Student’s T-test *p<0.05, **p < 0.005, ***p<0.0005. Graphs are means +/-SEM.

Delta insulin, delta AICAR and delta combination values showed a difference in pAS160 responses. There was an impaired DM1 pAS160 response after insulin treatment (Figure 7F). There were also trends (p = 0.0940) of impaired pAS160 response after AICAR treatment (Figure 7G). With combination treatment, we again observed a reduced DM1 pAS160 response (Figure 7H). Delta insulin and delta AICAR differences were lost after pAS160 was normalized to total AS160 (pAS160/AS160) (Figure 7I). Even after normalization, we still observed reduced DM1 pAS160/AS160 activation after combination treatment. Overall, we saw a consistent indication of impaired AS160 signaling in the DM1 myotubes (Figure 7L).

## 4. Discussion

### 4.1 Ingenuity Pathway Analysis of sedentary and exercised HSA-LR mice

This study was conducted to understand the insulin signaling pathway and exercise interaction within the context of DM1. Thus, we decided to investigate relevant targets pertaining to insulin and exercise signaling pathways. With IPA, we found some targets that aligned with the relevant pathways in sedentary and exercised HSA-LR mice. We decided to then acutely stimulate AMPK with AICAR in our HSA-LR mice and DM1 myotubes to determine the how AMPK-PGC-1α stimulation impacted insulin signaling. Our study, aligned with IPA findings, found that baseline total AMPK was lower in the HSA-LR mouse and in the DM1 myotubes. Additionally, in the HSA-LR mouse, PGC-1α expression did not respond to AICAR treatments as was found in WT mice, suggesting dysregulation of PGC-1α. Further, IPA predicted the insulin receptor to be inhibited in the sedentary HSA-LR mice. Our study found that pAkt signaling, a reporter of proximal insulin signaling, was indeed impaired in the HSA-LR mouse and the DM1 myotubes. After exercise intervention, IPA predicted PGC-1α and insulin receptor to be upregulated. However, in our DM1 models, we observed that acute AMPK stimulation did not improve PGC-1α or Akt levels or activity. Additionally, IPA detected increased GLUT4 expression in the exercised HSA-LR mice, indicating that exercise drives GLUT4 expression. Again, our findings did not align with IPA’s, as we saw no clear increase in total GLUT4 after AMPK-PGC-1α stimulation.

The powerful IPA analysis tool helped inform the investigation of targets pertinent to insulin and exercise signaling in DM1. The IPA analysis of the sedentary HSA-LR condition aligned with a lot of the findings we saw in the baseline expression of our DM1 models. However, when we decided to target the exercise axis, AMPK-PGC-1α, in an acute and targeted manner with AICAR, we saw that neither of the models had significant improvements in insulin resistance markers.

### 4.2 AICAR Treatment in HSA-LR Mouse Model

At baseline, total AMPK levels were lower in HSA-LR mice, however pAMPK was no different from WT. When subjected to AICAR, AMPK phosphorylation did not indicate any significant changes. Due to the transient nature of AMPK signaling, ACC phosphorylation was measured to validate the activity of the AICAR injection. Indeed, pACC increased, suggesting the increased activity of AMPK after AICAR treatment (Taylor et al., 2008).

Baseline expression of PGC-1α showed no differences between WT and HSA-LR. Yet, previous reports indicate a ∼43% lower expression of PGC-1α in HSA-LR skeletal muscle tissue (Ravel-Chapuis et al., 2018). Ravel-Chapuis et al. described this downregulation in extensor digitorum longus (EDL) muscle, whereas we analyzed protein content in the gastrocnemius. The HSA transgene is differentially expressed in muscles, with the number of mis-splicing events positively correlated with the expression of the HSA transgene across skeletal muscles (Hicks et al., 2024). It was found that the gastrocnemius had the highest abundance of HSA-LR construct expression which correlated with the highest mis-splicing events compared to the quadriceps and tibialis anterior muscles (Hicks et al., 2024). This research did not explicitly examine HSA construct expression in EDL muscle, but its findings may suggest why we found different results in gastrocnemius tissue.

After AICAR injection, PGC-1α levels did not change in HSA-LR mice, contrasting with the increase observed in WT mice. PGC-1α has a half-life ranging from 0.5-5.5 hours, as it is subjected to the ubiquitin protease degradation pathway (Trausch-Azar et al., 2010). AMPK drives the expression of PGC-1α and directly phosphorylates the transcription factor; however, AMPK has not been described to directly stabilize PGC-1α (Cantó and Auwerx, 2009a). Rather, p38 MAPK is implicated in phosphorylating PGC-1α and increasing its stability (Puigserver et al., 2001). Both exercise and AICAR have been reported to stimulate p38 activity, implicating the kinase as an intermediary within the AMPK-PGC-1α axis (Gibala et al., 2009; Lemieux et al., 2003). Additionally, activated AMPK upregulates transcription of PPARβ/δ, the receptor for PGC-1β and PGC-1δ, via the myocyte enhancer factor 2A (Koh et al., 2019). Then, PPARβ/δ stabilizes the PGC-1α expression levels in skeletal muscle tissue (Koh et al., 2017). The reduced baseline AMPK expression in HSA-LR mice may suggest that PGC-1α was not optimally regulated. Thus, in the acute treatment period of 30 minutes, we likely see an increased stability of the PGC-1α in WT mice which is reflected as elevated levels in western blotting.

Baseline GLUT4 levels were found to be downregulated in the HSA-LR mice. While this may not be explained by a difference in baseline PGC-1α levels, it could reflect diminished PGC-1α activity. For instance, phosphorylation and acetylation have both been shown to regulate PGC-1α transcriptional activity, which could alter expression levels of *SLC2A4* and the subsequent GLUT4 protein content (Michael et al., 2001; Wende et al., 2007a; Cantó and Auwerx, 2009a). There are no reports of *SLC2A4* mis-splicing in any DM1 models or patients, but mis-splicing could be a feasible explanation for the lower expression levels in HSA-LR tissues.

At baseline, there were no observed differences in pAkt, pAkt/Akt or pAS160 levels, but there were differences in their responses to AICAR. In WT mice, AICAR treatment increased pAkt and pAkt/Akt, similar to previous findings (Dedert et al., 2023). After repleting C2C12s with nutrient rich media, serine-473 pAkt increased via mTORC2 regulation (Dedert et al., 2023). This effect could explain why we observed an increase in pAkt for WT mice after AICAR treatment. While there is no conclusive literature that directly assesses Akt activity in HSA-LR mice, there is a study that explored GSK-3β, a downstream target inhibited by Akt (Lin et al., 2007). In line with our reports of impaired Akt activity, literature indicates that GSK-3β is upregulated in the HSA-LR mouse model (Jones et al., 2012). We also saw that pAS160 levels remained impaired and blunted even after AICAR injection in HSA-LR mice, which is consistent with the downregulated pAkt. In line with the impaired pAS160 signaling, we saw no significant change in the proportion of GLUT4 at the plasma membrane as evidenced by immunofluorescent staining. This contrasts with the response that we saw in WT mice, which had elevated levels of pAS160 and corresponding trends for increased plasma membrane-bound GLUT4. This is consistent with literature that finds AMPK stimulation, via exercise or AICAR, increases AS160 activity and GLUT4 shuttling (Arias et al., 2007b; Bradley et al., 2014; Wang et al., 2022). We observed that HSA-LR mice, compared to WT mice, consistently demonstrate a diminished response in Akt, PGC-1α, and pAS160. This outcome demonstrates the presence of insulin resistance in the HSA-LR mouse model and its resistance to acute AMPK stimulation.

### 4.3 Insulin and AICAR treatment in DM1 myotube models

Like the HSA-LR model, we saw downregulated expression of total AMPK in DM1 myotubes, however the pAMPK seemed to be upregulated. Given the transient nature of AMPK phosphorylation, we decided to assess pACC as stable readout of AMPK activity. We found proportionate phosphorylation of ACC after AICAR or combination treatment, indicating stimulation of AMPK. Contrary to what was observed in the HSA-LR skeletal muscle tissue, PGC-1α and GLUT4 did not show any differences at baseline or after stimulation with insulin or AICAR. This data is consistent with a study that found no differences in PGC-1α content before or after exercise intervention in DM1 patients (Mikhail et al., 2022).

The baseline serine-473 pAkt levels in DM1 myotubes were higher than healthy CTL myotube levels. Consistently, we saw an increased basal level of pAS160 in DM1 myotubes. Yet, following insulin and/or AICAR treatment, the DM1 myotubes had a lower delta response in pAkt, pAkt/Akt and pAS160. When Bertacca et al. subjected human myoblasts to a high concentration of insulin to induce resistance, they hyperactivated the insulin signaling pathway in the myoblasts (Bertacca et al., 2005). This hyperactivated insulin signaling pathway did not reflect higher glucose uptake at baseline and even had a blunted response to subsequent insulin treatment compared to healthy control myoblasts (Bertacca et al., 2005).This hyperactivated insulin signaling pathway did not reflect higher glucose uptake at baseline and even had a blunted response to subsequent insulin treatment compared to healthy control myoblasts (Bertacca et al., 2005). Their results mimic our findings, where elevated baseline Akt-AS160 signaling in our DM1 myotubes did not have CTL-like responses to insulin and/or AICAR treatment. AS160 phosphorylation leads to GLUT4 translocation, so our data may suggest that GLUT4 translocation and subsequent glucose uptake is impaired in the human DM1 myotubes.

### 4.4 Possible explanations for insulin resistance in DM1

The data from the HSA-LR mouse model and the DM1 myotube model share some similarities. In both models, we observe downregulated total AMPK. Despite this, we see that there is a proportional and adequate stimulation of pACC, indicating the stimulation of AMPK with AICAR. With the activation of AMPK in the HSA-LR model, we do not see an increase in PGC-1α protein. We did not see any changes in PGC-1α abundance in the DM1 myotube model, however this was no different than the CTL cells. Further, in both models we saw an impaired response to treatment in Akt phosphorylation. Consistently, reduced AS160 phosphorylation was observed after AICAR treatment and following insulin treatment in the myotube model. While we see these differences shared across both models, there is no literature describing the mis-splicing of RNA transcripts for AMPK, PGC-1α, Akt, AS160 or GLUT4. *INSR* can be mis-spliced in DM1, but this mechanism cannot account for all our findings since *INSR* is not mis-spliced in the HSA-LR model (Brockhoff et al., 2017). However, there is the prediction that a protein implicated in insulin signaling, Tre-2/Bub2/Cdc16 (TBC) domain family member 15 (TBC1D15), may be dysregulated because of RNA mis-splicing (Nakamori et al., 2013).

TBC1D15 is specifically implicated in the Rab protein interaction with GSVs. Rab GTPase proteins function to transduce the signaling to mechanical effectors like myosin and actin to initiate the movement of GSVs (Klip et al., 2014). TBC1D15 then serves as GAP, similar to AS160, for Rab GTPase proteins (Peralta et al., 2010). TBC1D15 is also important for regulating lysosome and mitochondrial morphology (Yamano et al., 2014). Wu et al. knocked TBC1D15 out of different cell lines including the L6 rate skeletal muscle cell line and found that GLUT4 content was reduced (Wu et al., 2019). Interestingly, one group predicts that TBC1D15 may be mis-spliced in DM1 (Nakamori et al., 2013), which could possibly explain the post-insulin receptor dysregulations found in our study. This could also provide further context as to why GLUT4 content was reduced in the HSA-LR tissue. Validating TBC1D15 at the RNA and protein levels in future studies would help further explore the dysregulated insulin signaling pathway in DM1.

### 4.5 Significance

Insulin signaling is key to the anabolic processes that occur in the cell to advance cell growth and cell division. Therefore, it is important to investigate insulin resistance in DM1, as it can exacerbate some of the other symptoms implicated in the disorder such as muscle atrophy and weakness (Renna et al., 2019). Studies have assessed the use of classical insulin sensitizing treatments, like metformin, in DM1, however the impacts of these treatments on insulin resistance in DM1 were not extensively explored (Gagnon et al., 2018; García-Puga et al., 2022). Our study sought to provide context on the response for insulin signaling molecules in DM1 following acute AMPK stimulation. In doing so, we were able to explore the responses in two models of DM1 where we confirmed dysregulated insulin signaling, prior to and after AMPK stimulation.

Exercise yields improvements in strength, mobility and metabolism for DM1 patients, but insulin resistance has not been specifically assessed (Okkersen et al., 2018; Roussel et al., 2019, 2020; Di Leo et al., 2023). Metabolic adaptations from exercise are classically linked with improvements in glucose uptake which necessitates the investigation of the insulin signaling pathway. However, other factors such as elevated local blood flow and blood glucose concentration can also contribute to exercised-induced glucose uptake (Martin et al., 1995; Rose and Richter, 2005). This study has demonstrated the downregulated behaviour of insulin signaling molecules in response to AMPK stimulation in DM1 but does not discount the overall validity of exercise for DM1 patients. The design of our study specifically assessed acute AMPK stimulation, which means that DM1 patients could still see insulin signaling benefits with chronic AMPK stimulation.

### 4.6 Limitations

Key to this study was the stimulation of the exercise pathway through the AMPK-PGC-1α axis using acute AICAR treatment. We decided to study this in a female HSA-LR mice along with cells derived from patients that were not sex-matched. However, we recognize that DM1 can present in a sex-specific manner, with further sex-specific responses to exercise (Dogan et al., 2016; Ravel-Chapuis et al., 2018). In future studies, it would be ideal to sex-match pre-clinical models to gather a clear scope for sex-specific modifiers.

Due to the transient nature of the AMPK and AS160 phosphorylation, mice were treated for 30 minutes with AICAR, while myotubes were treated with it for 60 minutes (Sriwijitkamol et al., 2007; Funai and Cartee, 2008). It would be insightful to explore time course treatments to investigate the interactions of the proteins after chronic AMPK stimulation. Further, investigations with acute and chronic exercise regimens in the mice could elucidate some of the long term and systemic impacts that exercise has on the insulin signaling pathway.

### 4.7 Conclusions

After employing both a mouse and human myotube model, this study demonstrated that the insulin signaling pathway is impaired in DM1 and is not rescued with acute AMPK stimulation. Namely, we found that acute AMPK activation did not rescue the Akt-AS160 signaling pathway in either of the models, suggesting that GLUT4 translocation is impaired in DM1 prior to and after AMPK activation. Further studies could confirm RNA levels of the insulin and exercise pathway proteins to investigate possible spliceopathies. A target of note is TBC1D15, which is involved in GLUT4 stability and translocation and is predicted to be mis-spliced in DM1. Further, time course treatments with AICAR, and long-term exercise studies in mice and humans could further elucidate the interactions of the insulin signaling and exercise signaling proteins. Altogether, this study provides new insights into the pathomechanisms of insulin resistance in DM1 and furthers the understanding of metabolic behaviour in DM1 patients.

## Supporting information

Supplemental Excel Sheets

## Acknowledgments

We would like to extend thanks to Dr. Elena Pegoraro’s group and patient donors for granting us with the patient-derived myoblasts that were implemented in this study.

## Contributions

O.A. planned and executed experimentation detailed in the study. He analyzed data, wrote and revised manuscript.

S.S. managed resource allocation, provided guidance and insight for experimentation. Edited and revised manuscript.

R.C. helped with mouse injections and dissections. He also helped prepare cryosections and analysis workflow for immunofluorescent imaging. Also helped edit manuscript.

A.M conducted original exercise study on HSA-LR mice which were used in proteomic analysis. Provided directional insight for study scope and aims.

A.H conducted mass spectrometry analysis on sedentary and exercised HSA-LR mice.

K.H., D.O. helped with mouse injections and dissections. Also helped edit manuscript.

L.D. managed mice in housing facility and assisted with mouse injections and dissections.

A.R., V.L. managed and oversaw sedentary and exercised HSA-LR tissues were processed for mass spectrometry.

A.R.C. was an experimental advisor for the study. Provided HSA-LR mice used in AICAR study. Provided edits and revisions to writing.

A.M. was an experimental advisor for the study. Provided edits and revisions to writing.

H.L. Principal investigator of the project, providing guidance, resources, advice for experimental direction. Helped edits and revisions to writing.

