## Supplemental Document for "Acute AMPK activation does not adequately stimulate insulin signaling in skeletal muscle models of Myotonic Dystrophy Type 1"

1.

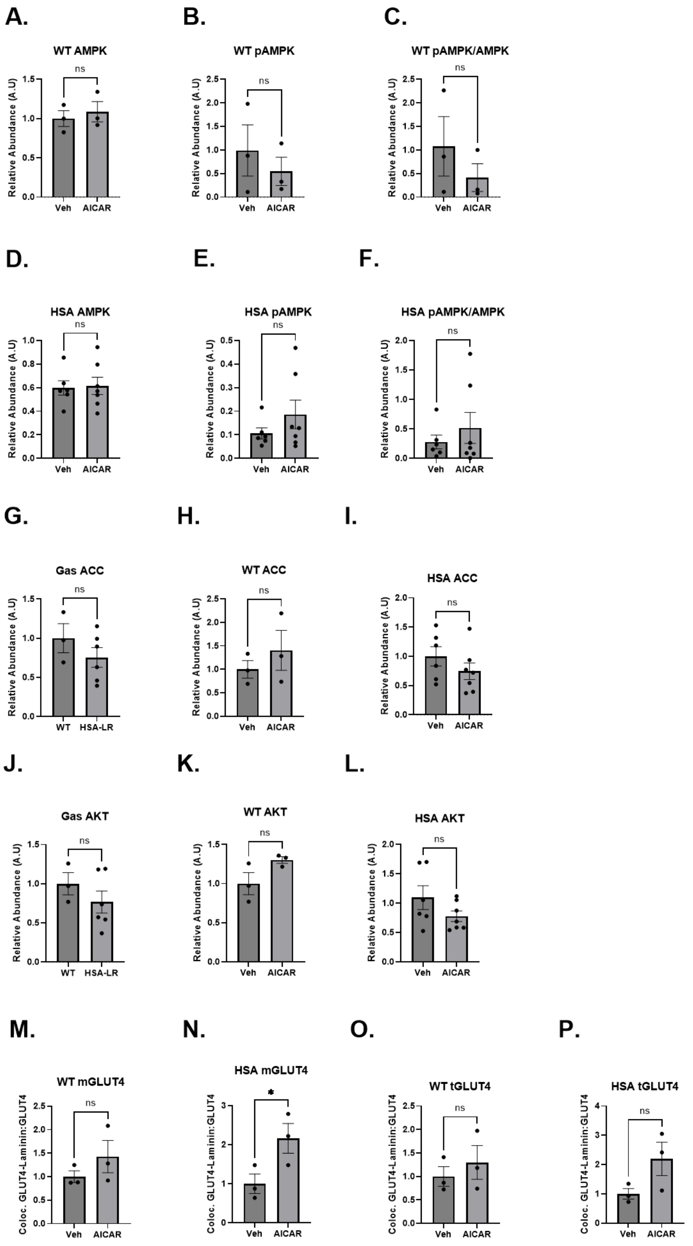

Supplemental Figure 1. Mouse Western blot and IF Quantifications. A.-C. WT comparison of A) total AMPK, B) pAMPK, and C) pAMPK/AMPK before and after AICAR treatment. D.-F. HSA-LR comparison of D) total AMPK, E) pAMPK, and F) pAMPK/AMPK before and after AICAR treatment. G.-I. Total ACC comparison at G) baseline, after AICAR treatment in H) WT and I) HSA-LR mice. J.-L. Total AKT comparison at J) baseline, after AICAR in K) WT and L) HSA-LR mice. M.-P. membrane bound GLUT4 (mGLUT4) response after AICAR treatment in

M) WT and N) HSA-LR and total GLUT4 (tGLUT4) response after AICAR treatment in O) WT and P) HSA-LR mice.

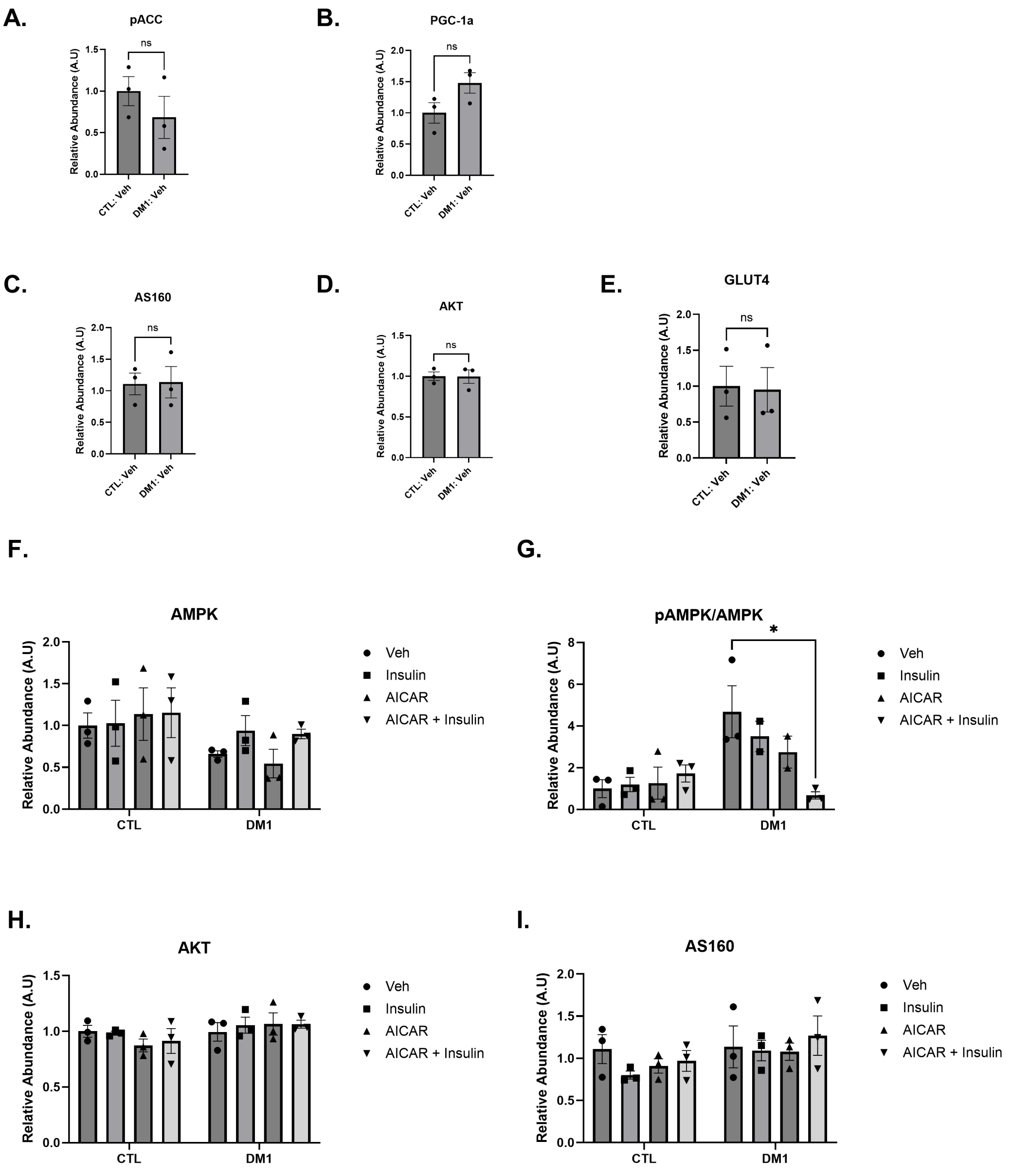

Supplemental Figure 2. Myotube Western blot Quantifications. A.-E. No difference in baseline A. pACC, B. PGC-1a, C. AS160, D. AKT, or E. GLUT4 in control (CTL) or myotonic dystrophy type 1 (DM1) myotubes.

F. No change in total adenosine monophosphate activated protein kinase (AMPK) levels after insulin, AICAR or combination treatment in CTL or DM1 myotubes. G. Decreased pAMPK/AMPK detected after combination treatment in DM1 myotubes. H., I. No difference detected in total H) AKT or I) AS160 levels across treatment groups or conditions in in CTL or DM1 myotubes.

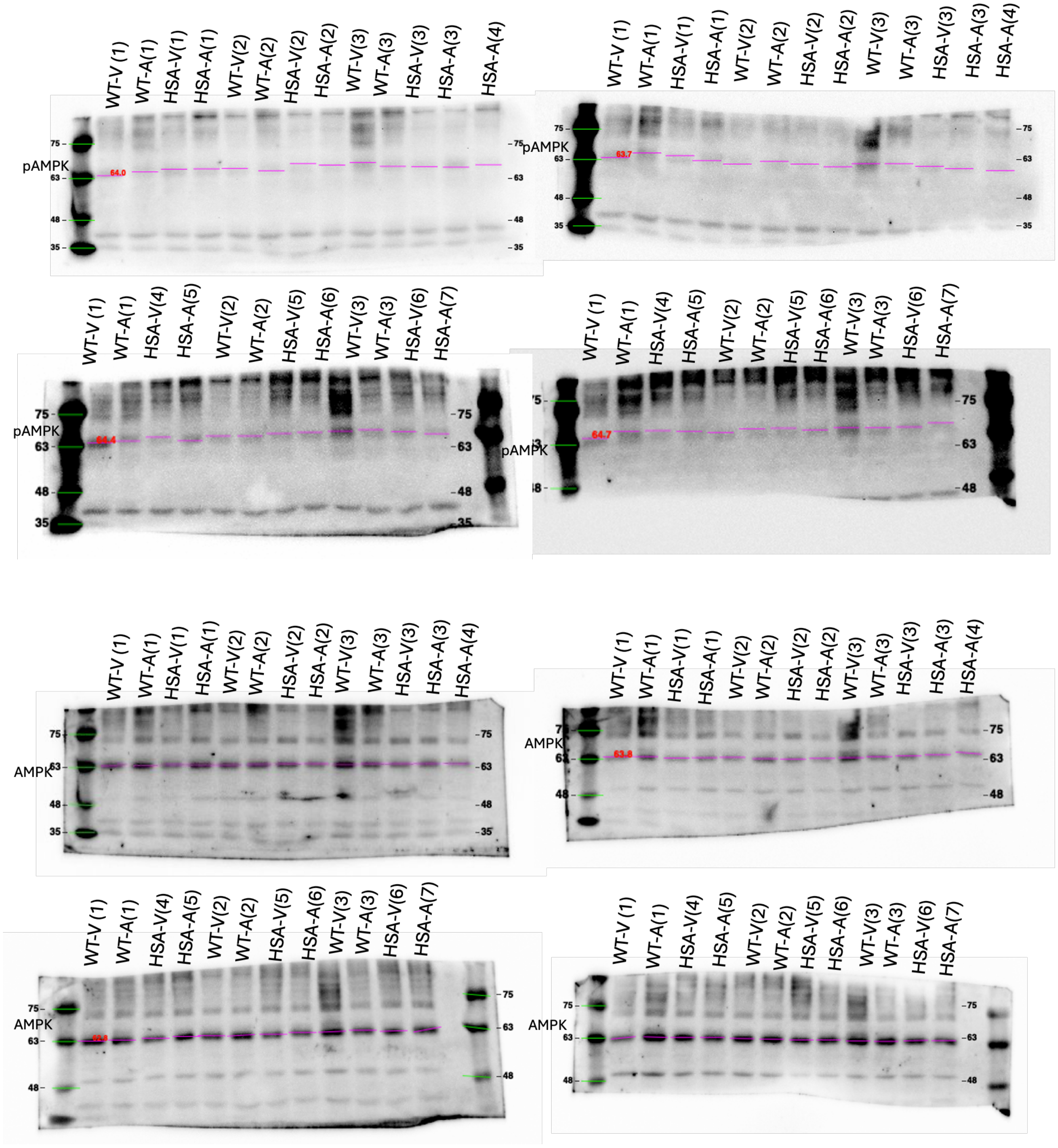

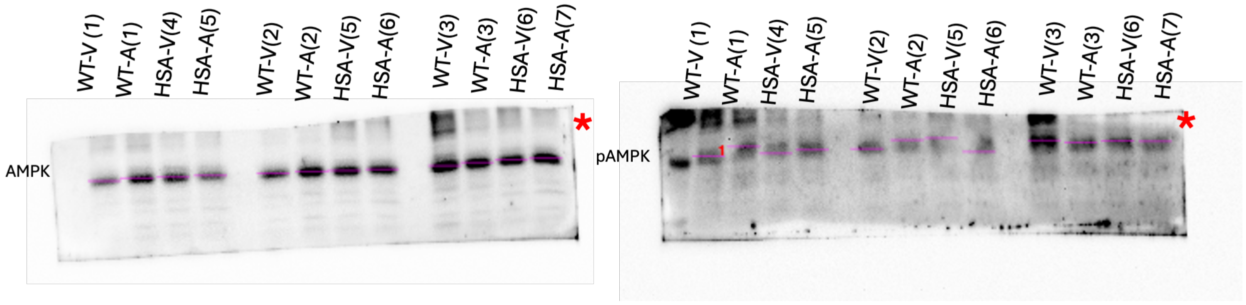

Supplemental Figure 3. Uncropped total and phosphorylated (Threonine 172) AMPK mouse western blots for Figure 2A. Expected banding at 62 kDa. Western blot banding found above ~64 kDa (Pink lines). Red asterisks indicates the blot used for representative blot. “WT” represents wildtype mice and “HSA” represents human skeletal actin long repeat mice. “V” represents vehicle treatment and “A” represents AICAR treatment.

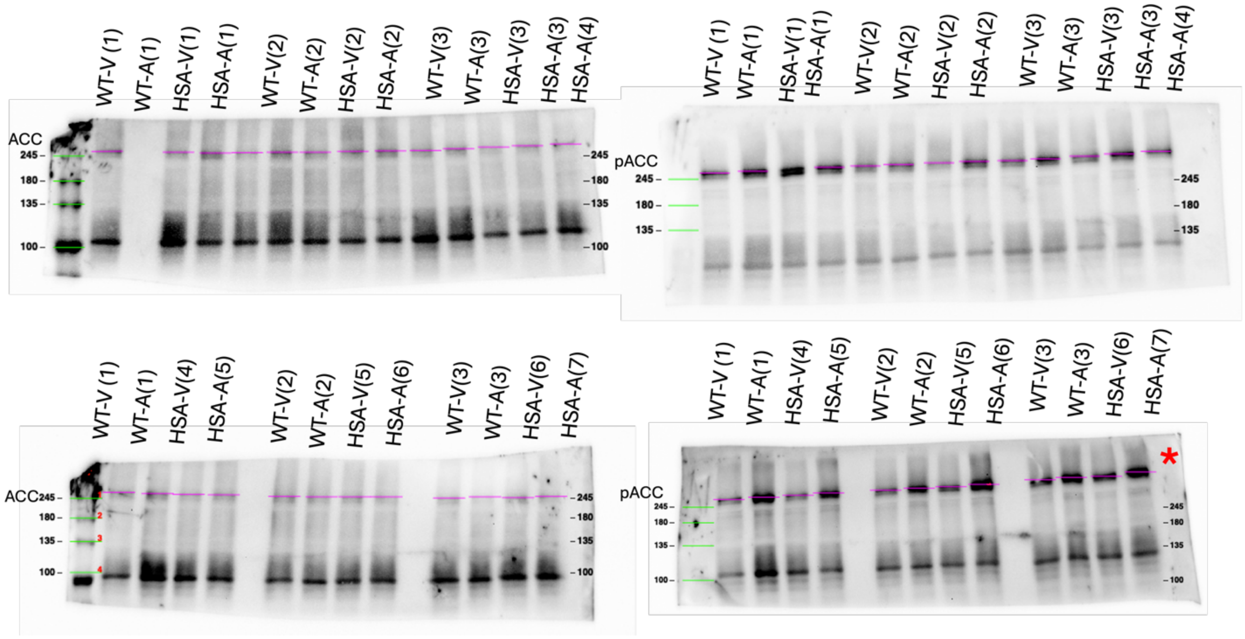

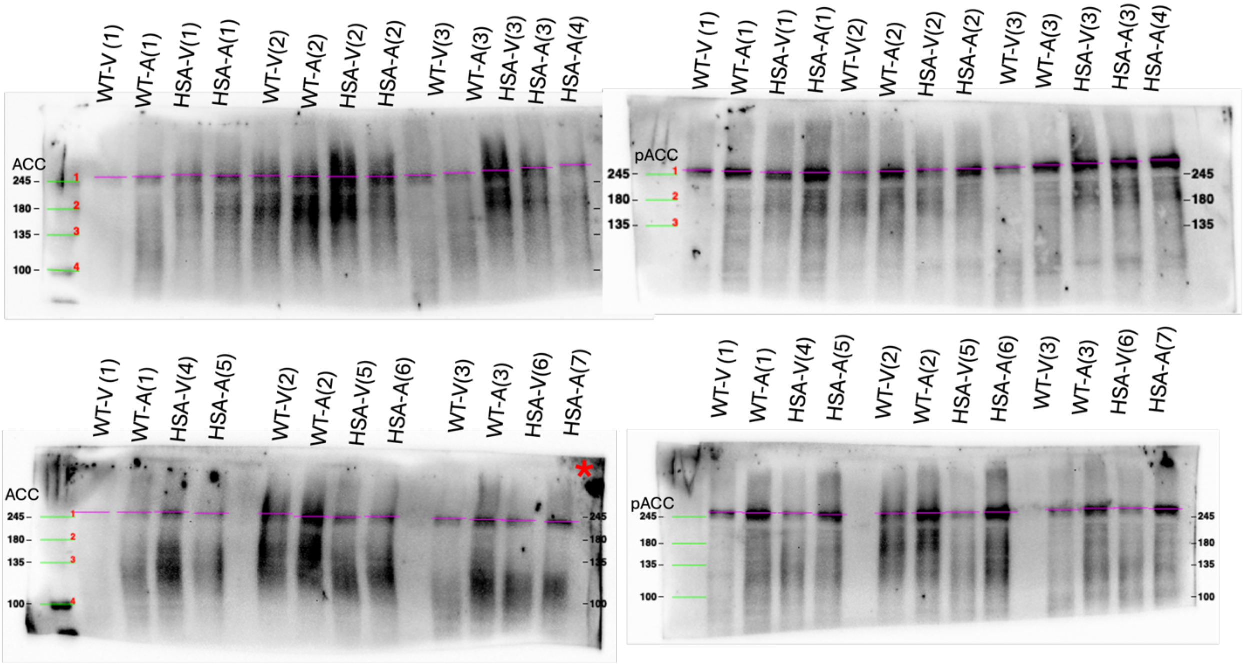

Supplemental Figure 4. Uncropped total and phosphorylated (Serine 79) ACC mouse western blots for Figure 2A. Expected banding at 280 kDa. Western blot banding found above ~245 kDa (Pink lines). Red asterisks indicates the blot used for representative blot. “WT” represents wildtype mice and “HSA” represents human skeletal actin long repeat mice. “V” represents vehicle treatment and “A” represents AICAR treatment.

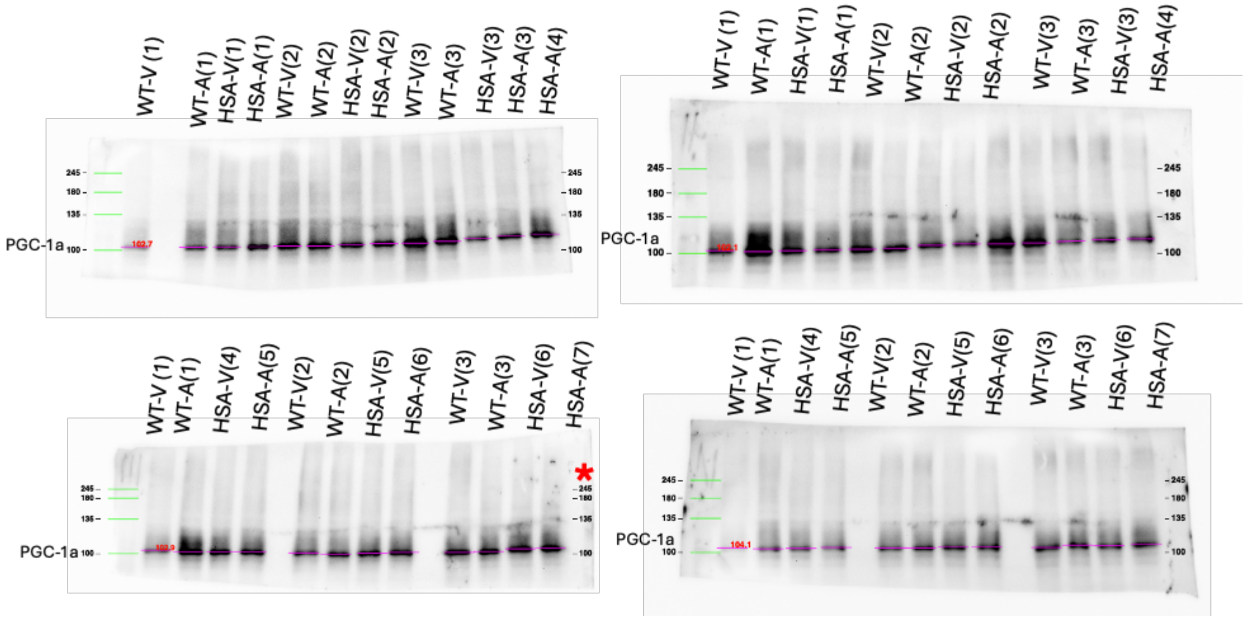

Supplemental Figure 5. Uncropped total PGC-1a mouse western blots used for Figure 2A. Expected banding between 90-100 kDa. Western blot banding found at ~100-105 kDa (Pink lines). Red asterisks indicates the blot used for representative blot. “WT” represents wildtype mice and “HSA” represents human skeletal actin long repeat mice. “V” represents vehicle treatment and “A” represents AICAR treatment.

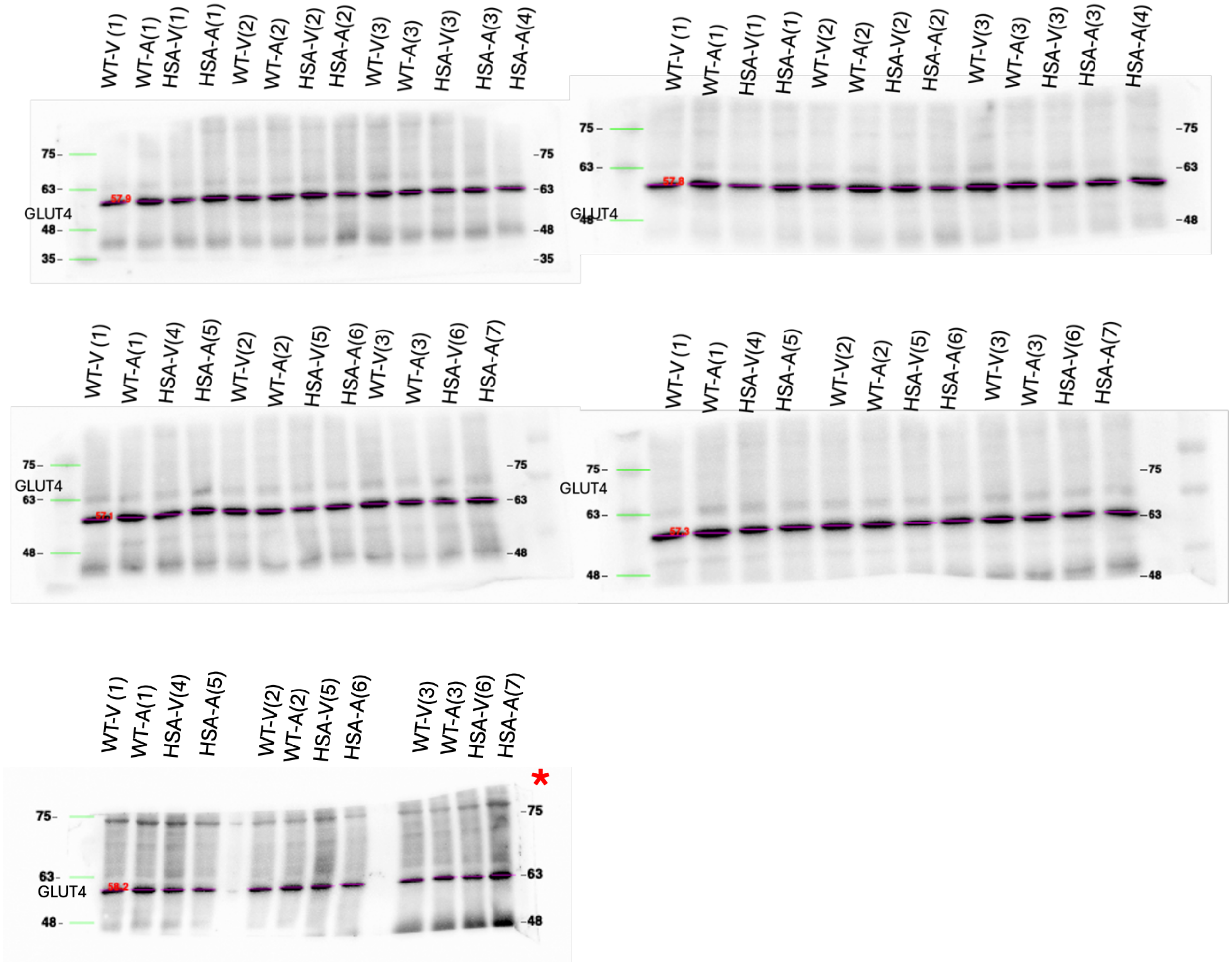

Supplemental Figure 6. Uncropped total GLUT4 mouse western blots used for Figure 2A. Expected banding between ~50-55kDa. Western blot banding found at ~55-60 kDa (Pink lines). Red asterisks indicates the blot used for representative blot. “WT” represents wildtype mice and “HSA” represents human skeletal actin long repeat mice. “V” represents vehicle treatment and “A” represents AICAR treatment.

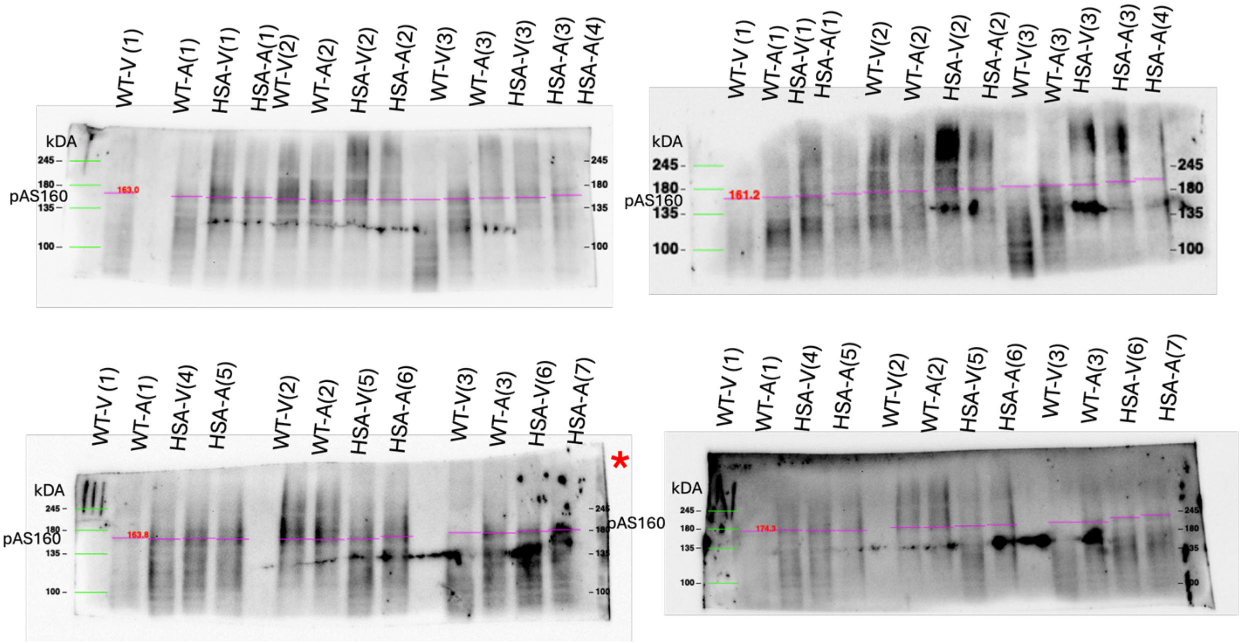

Supplemental Figure 7. Uncropped phosphorylated (Threonine 642) AS160 blots used for Figure 3A. Expected banding at 160 kDa. Western blot banding found between ~160-175 kDa (Pink lines). Red asterisks indicate the blot used for representative blot. "WT" represents wildtype mice and "HSA" represents human skeletal actin long repeat mice. "V" represents vehicle treatment and "A" represents AICAR treatment.

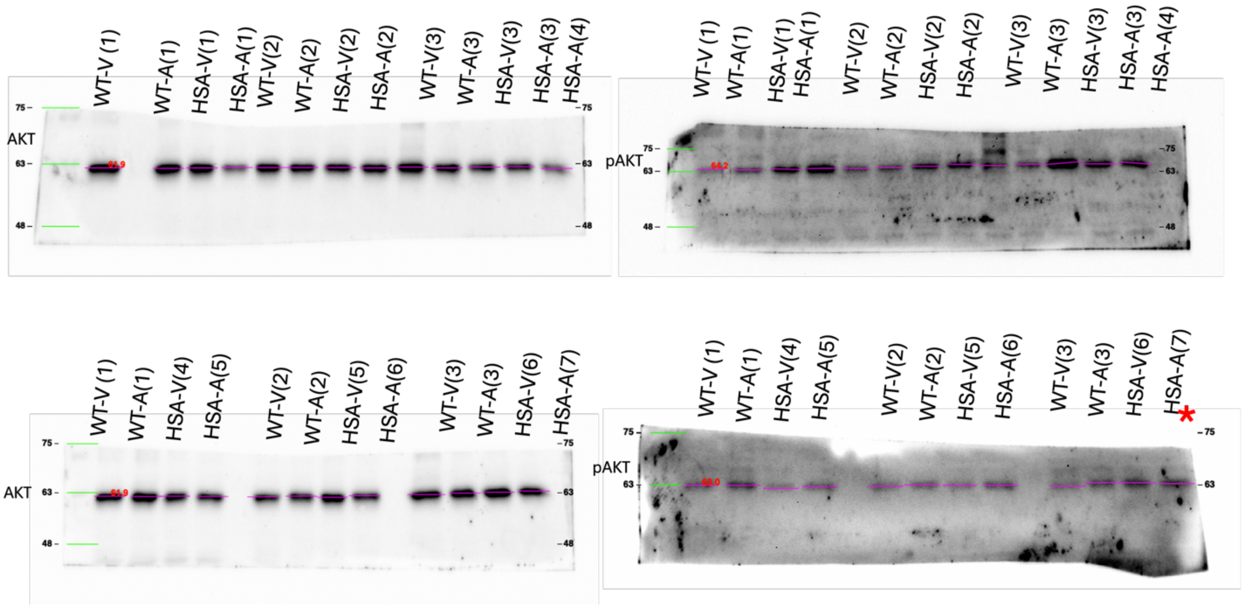

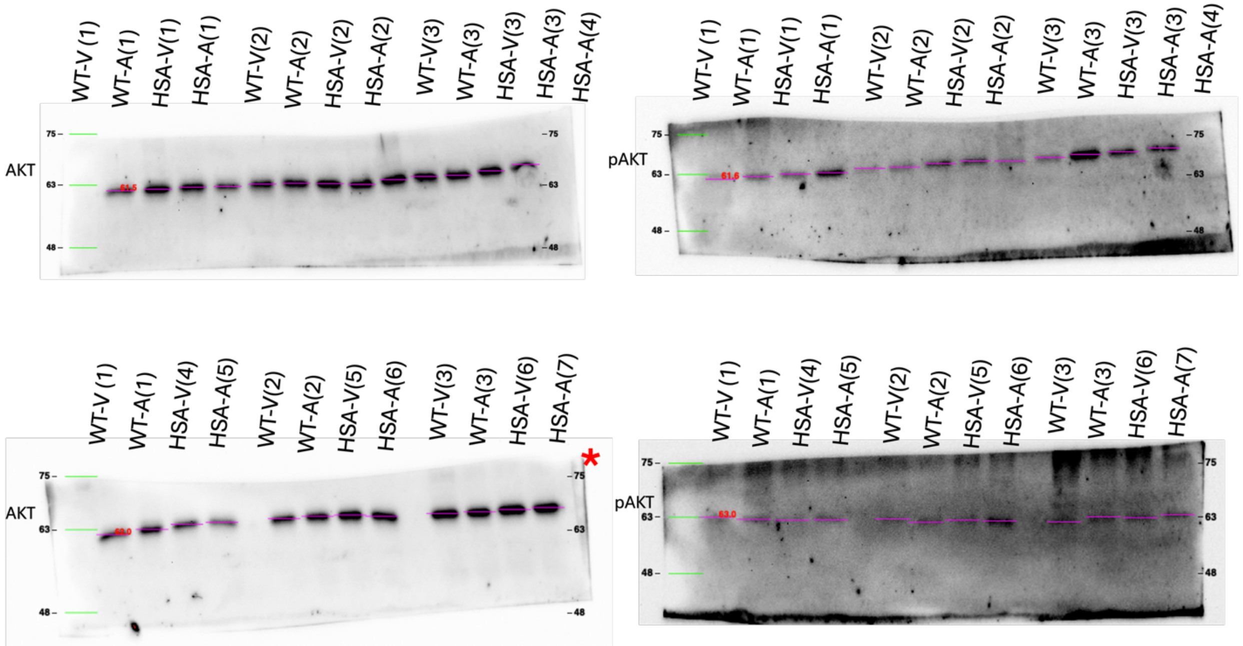

Supplemental Figure 8. Uncropped total and phosphorylated (Serine 473) AKT mouse western blots used for Figure 3A. Expected banding at 60 kDa. Western blot banding found between ~61-64 kDa (Pink lines). Red asterisks indicates the blot used for representative blot. “WT” represents wildtype mice and “HAS” represents human skeletal actin long repeat mice. “V” represents vehicle treatment and “A” represents AICAR treatment.

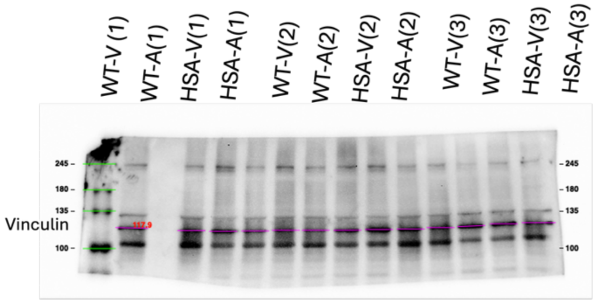
WT-V (1)
WT-A(1)
HSA-V(1)
HSA-A(1)
WT-V(2)
WT-A(2)
HSA-V(2)
HSA-A(2)
WT-V(3)
WT-A(3)
HSA-V(3)
HSA-A(3)
HSA-A(4)
Vinculin
245
180
135
100

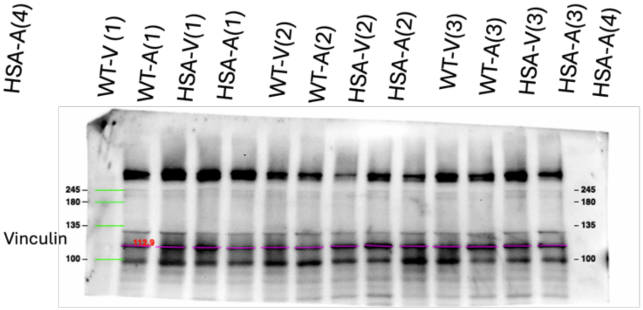
WT-V (1)
WT-A(1)
HSA-V(1)
HSA-A(1)
WT-V(2)
WT-A(2)
HSA-V(2)
HSA-A(2)
WT-V(3)
WT-A(3)
HSA-V(3)
HSA-A(3)
HSA-A(4)
Vinculin
245
180
135
100

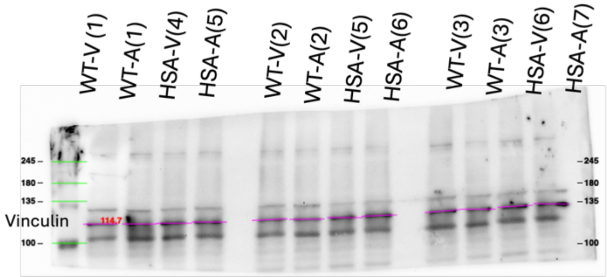
WT-V (1)
WT-A(1)
HSA-V(4)
HSA-A(5)
WT-V(2)
WT-A(2)
HSA-V(5)
HSA-A(6)
WT-V(3)
WT-A(3)
HSA-V(6)
HSA-A(7)
Vinculin
245
180
135
100

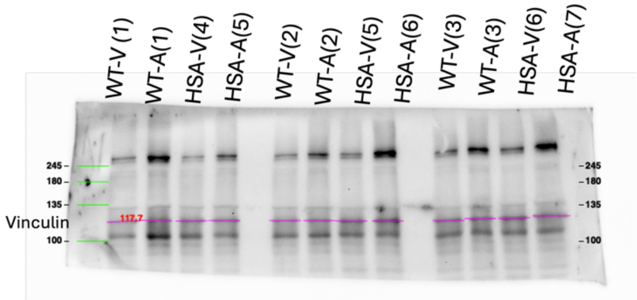
WT-V (1)
WT-A(1)
HSA-V(4)
HSA-A(5)
WT-V(2)
WT-A(2)
HSA-V(5)
HSA-A(6)
WT-V(3)
WT-A(3)
HSA-V(6)
HSA-A(7)
Vinculin
245
180
135
100

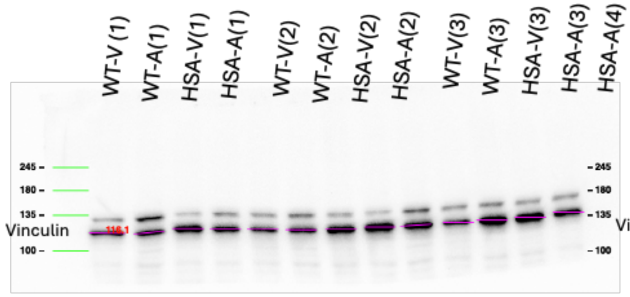
WT-V (1)
WT-A(1)
HSA-V(1)
HSA-A(1)
WT-V(2)
WT-A(2)
HSA-V(2)
HSA-A(2)
WT-V(3)
WT-A(3)
HSA-V(3)
HSA-A(3)
HSA-A(4)
Vinculin
245
180
135
100

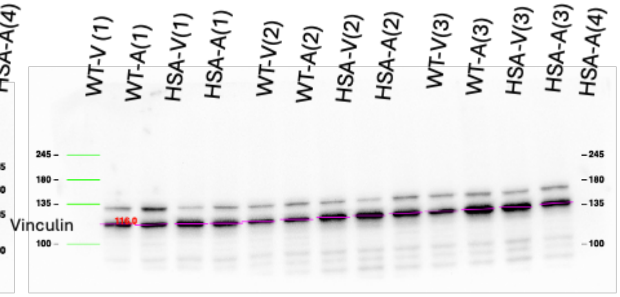
WT-V (1)
WT-A(1)
HSA-V(1)
HSA-A(1)
WT-V(2)
WT-A(2)
HSA-V(2)
HSA-A(2)
WT-V(3)
WT-A(3)
HSA-V(3)
HSA-A(3)
HSA-A(4)
Vinculin
245
180
135
100

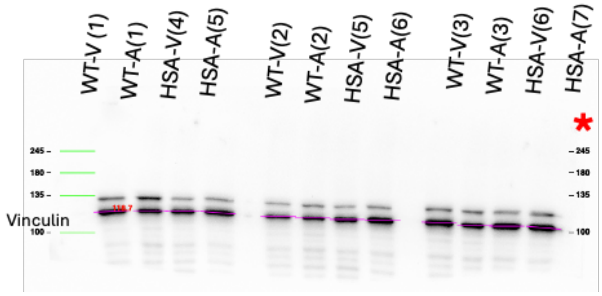
WT-V (1)
WT-A(1)
HSA-V(4)
HSA-A(5)
WT-V(2)
WT-A(2)
HSA-V(5)
HSA-A(6)
WT-V(3)
WT-A(3)
HSA-V(6)
HSA-A(7)
*
Vinculin
245
180
135
100

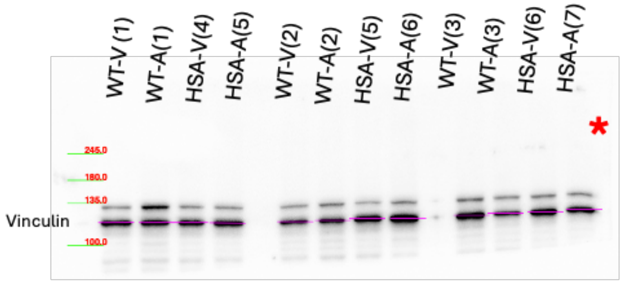
WT-V (1)
WT-A(1)
HSA-V(4)
HSA-A(5)
WT-V(2)
WT-A(2)
HSA-V(5)
HSA-A(6)
WT-V(3)
WT-A(3)
HSA-V(6)
HSA-A(7)
*
Vinculin
245.0
180.0
135.0
100.0

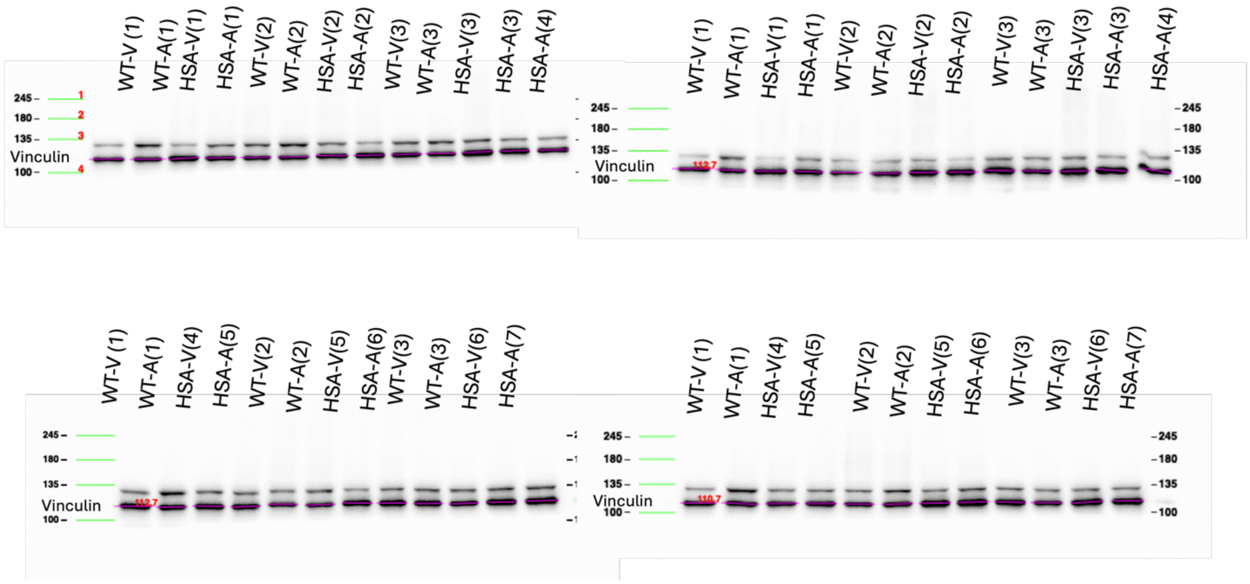

Supplemental Figure 9. Uncropped total vinculin mouse western blots used for Figure 2A and 3A. Expected banding at ~124 kDa. Western blot banding found at ~110-120 kDa (Pink lines). Red asterisks indicates the blot used for representative blot. “WT” represents wildtype mice and “HSA” represents human skeletal actin long repeat mice. “V” represents vehicle treatment and “A” represents AICAR treatment.

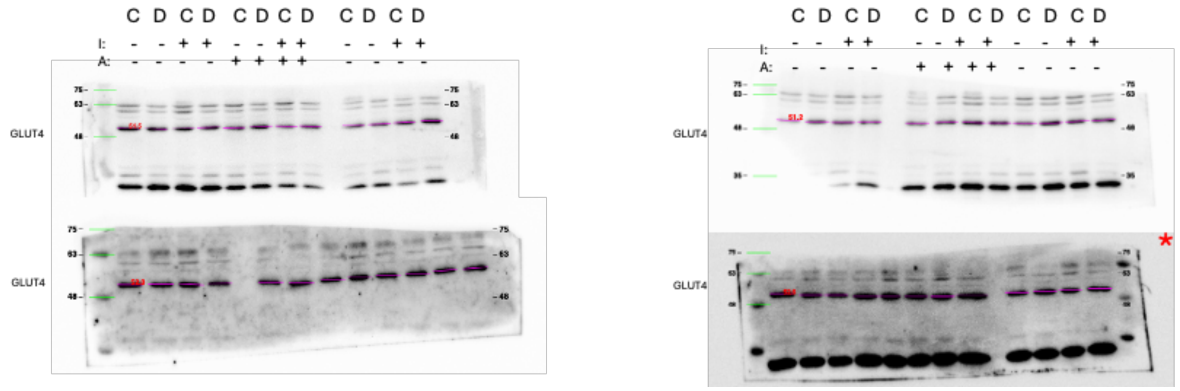

Supplemental Figure 10. Uncropped total GLUT4 myotube western blots used for Figure 5A. Expected banding at 50-55 kDa. Western blot banding found between ~50-55 kDa (Pink lines). Red asterisks indicates the blot used for representative blot. “C” represents Control myotubes and “D” represents DM1 myotubes. “I” represents insulin treatment and “A” represents AICAR treatment. ”nc” represents normalizing control.

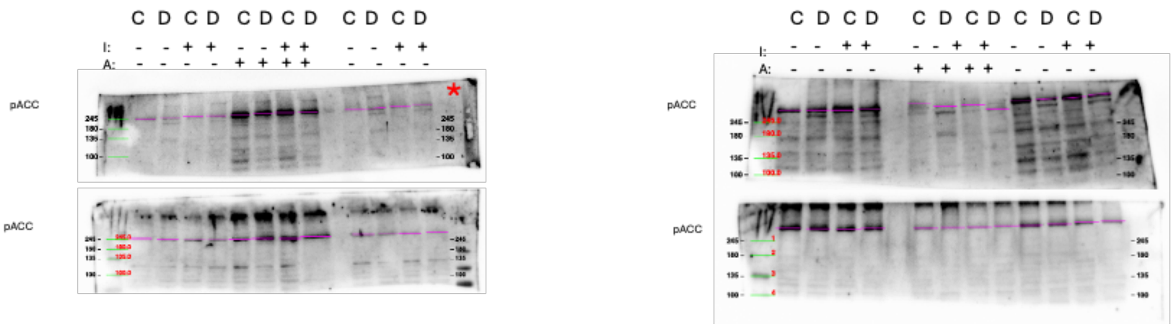

Supplemental Figure 11. Uncropped phosphorylated (Serine 79) ACC myotube western blots for Figure 5A. Expected banding at ~285 kDa. Western blot banding above ~245 kDa (Pink lines). Red asterisks indicates the blot used for representative blot. “C” represents Control myotubes and “D” represents DM1 myotubes. “I” represents insulin treatment and “A” represents AICAR treatment. ”nc” represents normalizing control.

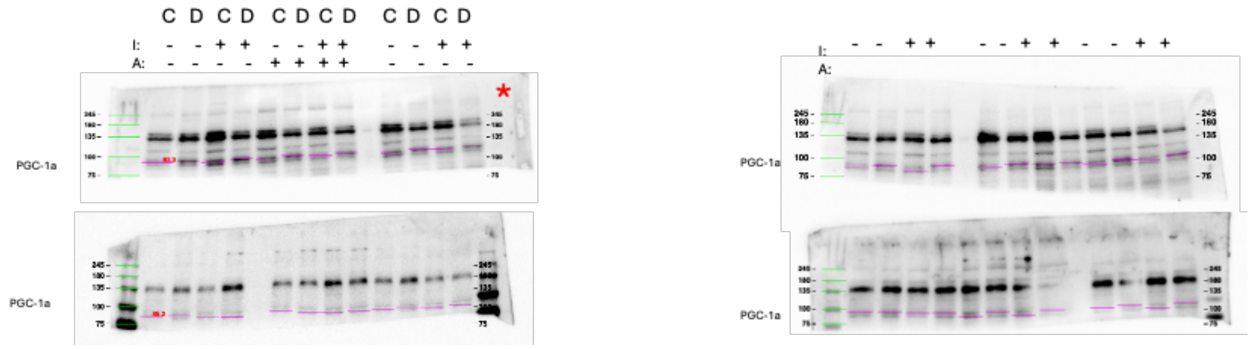

Supplemental Figure 12. Uncropped total PGC-1a myotube western blots for Figure 5A. Expected banding at 90-100 kDa. Western blot banding found between ~85-95 kDa (Pink lines). Red asterisks indicates the blot used for representative blot. “C” represents Control myotubes and “D” represents DM1 myotubes. “I” represents insulin treatment and “A” represents AICAR treatment. ”nc” represents normalizing control.

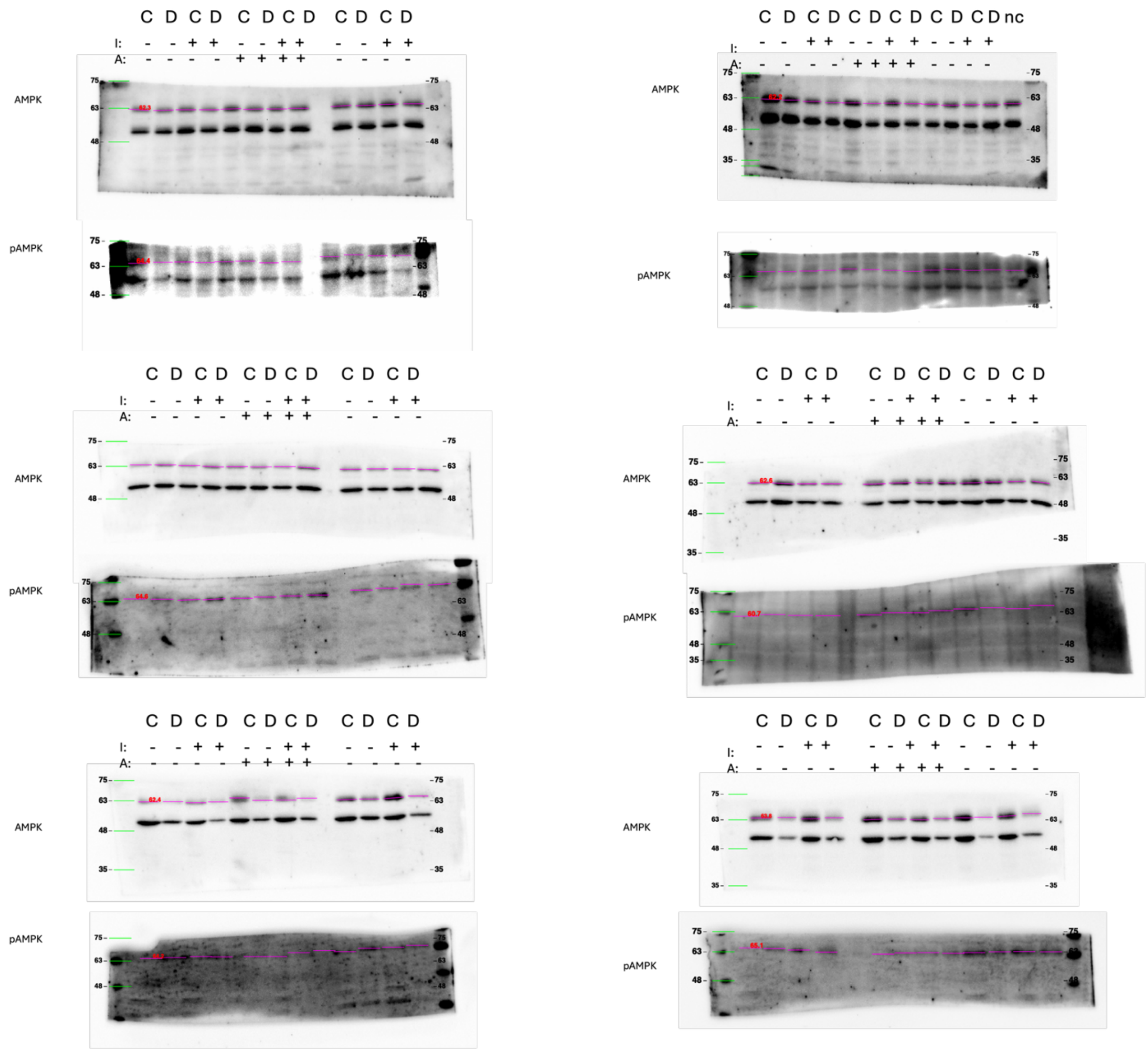

Supplemental Figure 13. Uncropped total and phosphorylated (Threonine 172) AMPK myotube western blots used for Figure 5A. Expected banding at 62 kDa. Western blot banding found between ~60-65 kDa (Pink lines). Red asterisks indicates the blot used for representative blot. “C” represents Control myotubes and “D” represents DM1 myotubes. “I” represents insulin treatment and “A” represents AICAR treatment. ”nc” represents normalizing control.

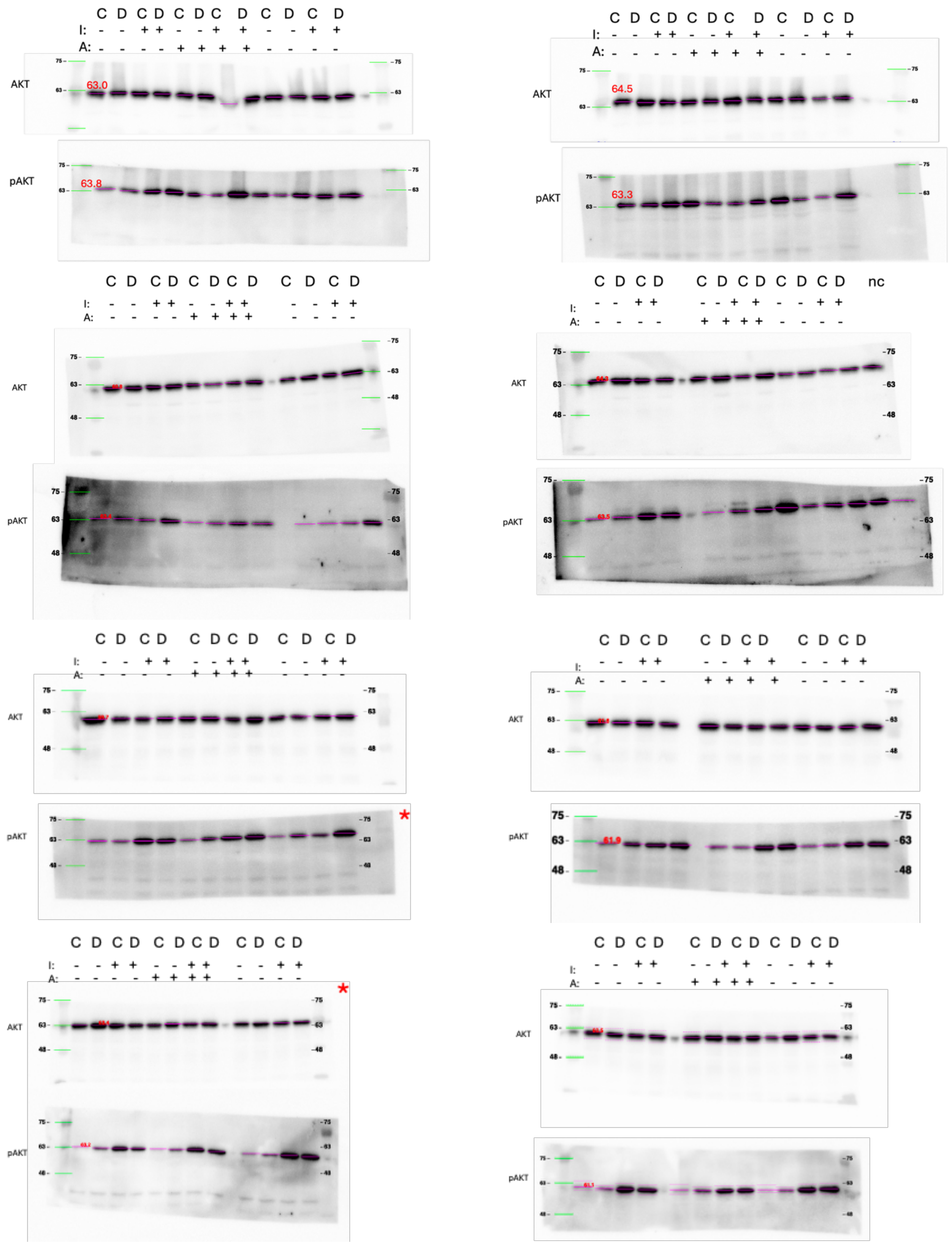

C D C D C D C D C D C D
I: - - + + - - + + - - + +
A: - - - - + + + + - - - -
AKT
pAKT
63.0
63.8
64.5
63.3
nc
75
63
48

Supplemental Figure 14. Uncropped total and phosphorylated (Serine 473) AKT myotube western blots used for Figure 6A. Expected banding at 60 kDa. Western blot banding found between ~60-65 kDa (Pink lines). Red asterisks indicates the blot used for representative blot. "C" represents Control myotubes and "D" represents DM1 myotubes. "I" represents insulin treatment and "A" represents AICAR treatment. "nc" represents normalizing control.

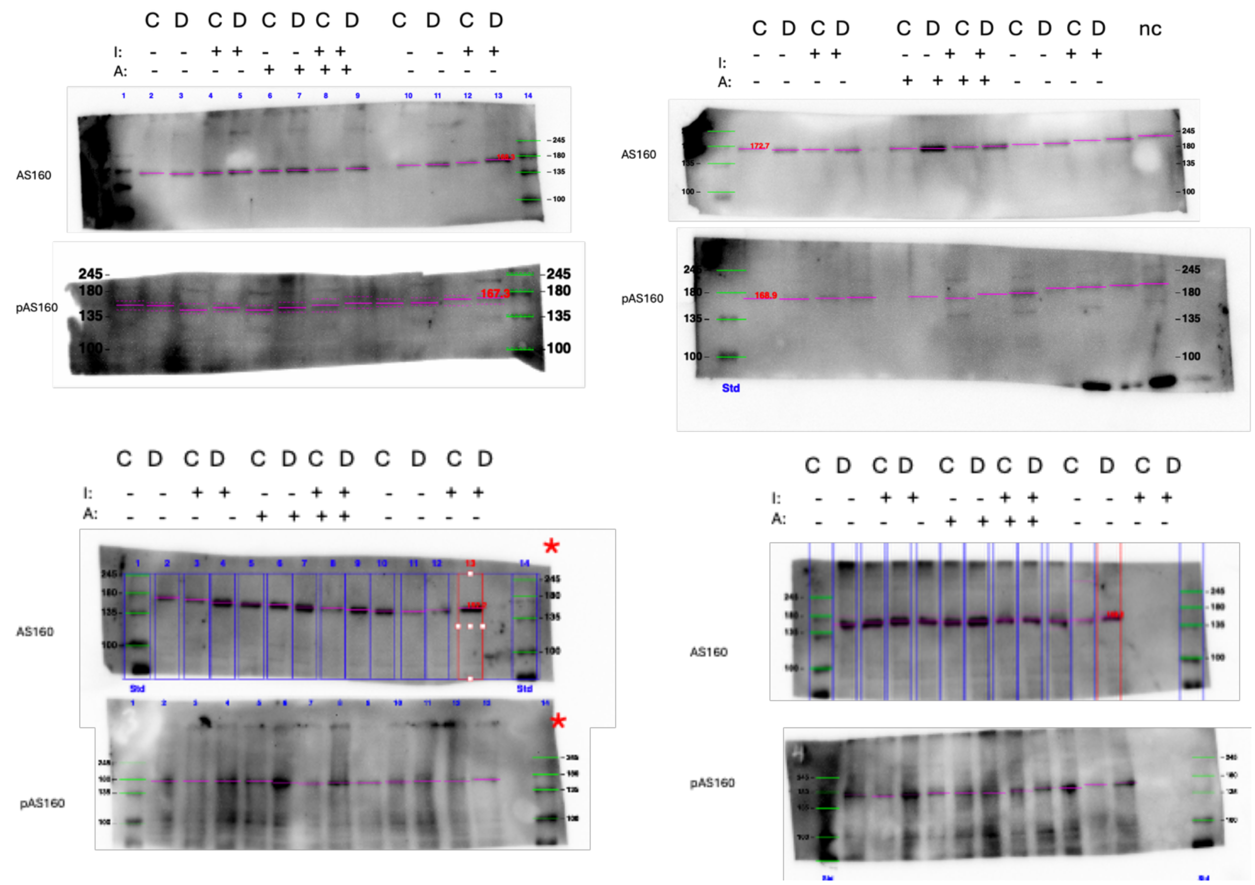

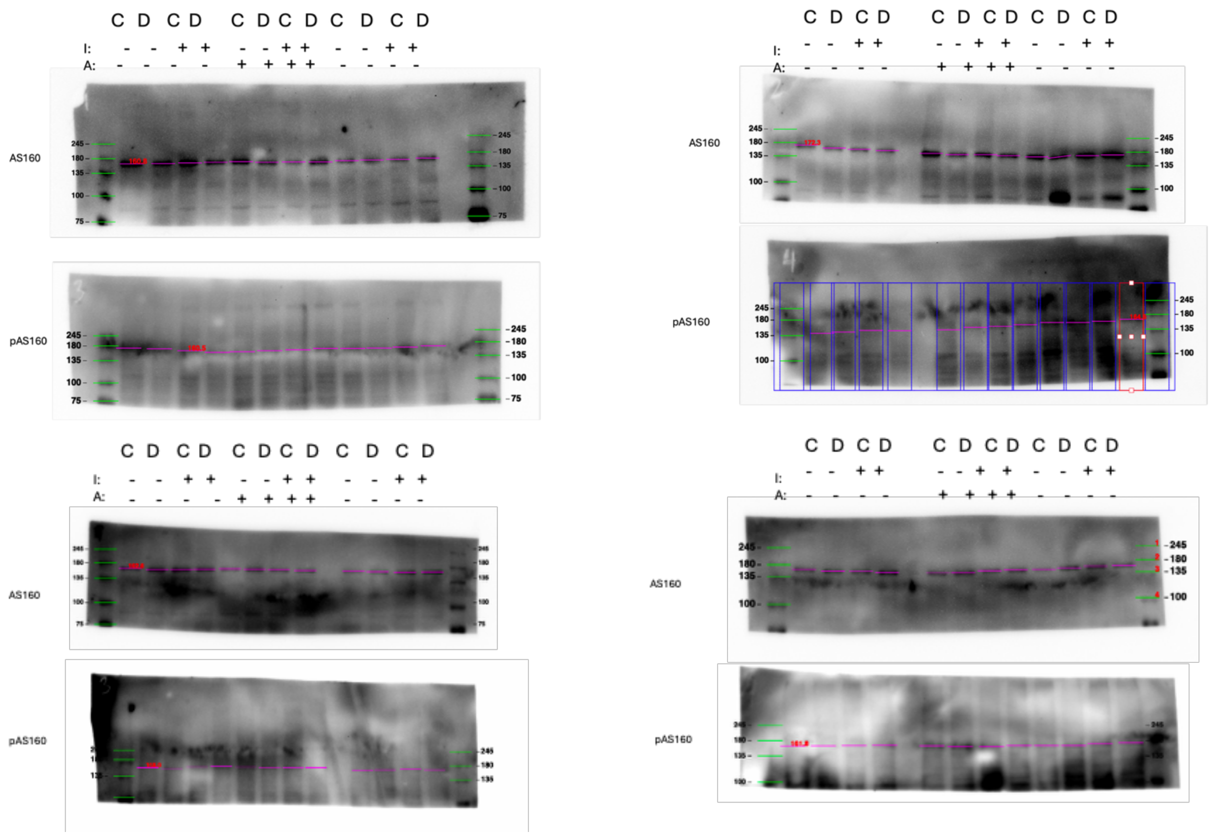

Supplemental Figure 15. Uncropped total and phosphorylated (Threonine 642) AS160 myotube western blots used for Figure 7A. Expected banding at 160 kDa. Western blot banding found between ~160-175 kDa (Pink lines). Red asterisks indicates the blot used for representative blot. "C" represents Control myotubes and "D" represents DM1 myotubes. "I" represents insulin treatment and "A" represents AICAR treatment. "nc" represents normalizing control.

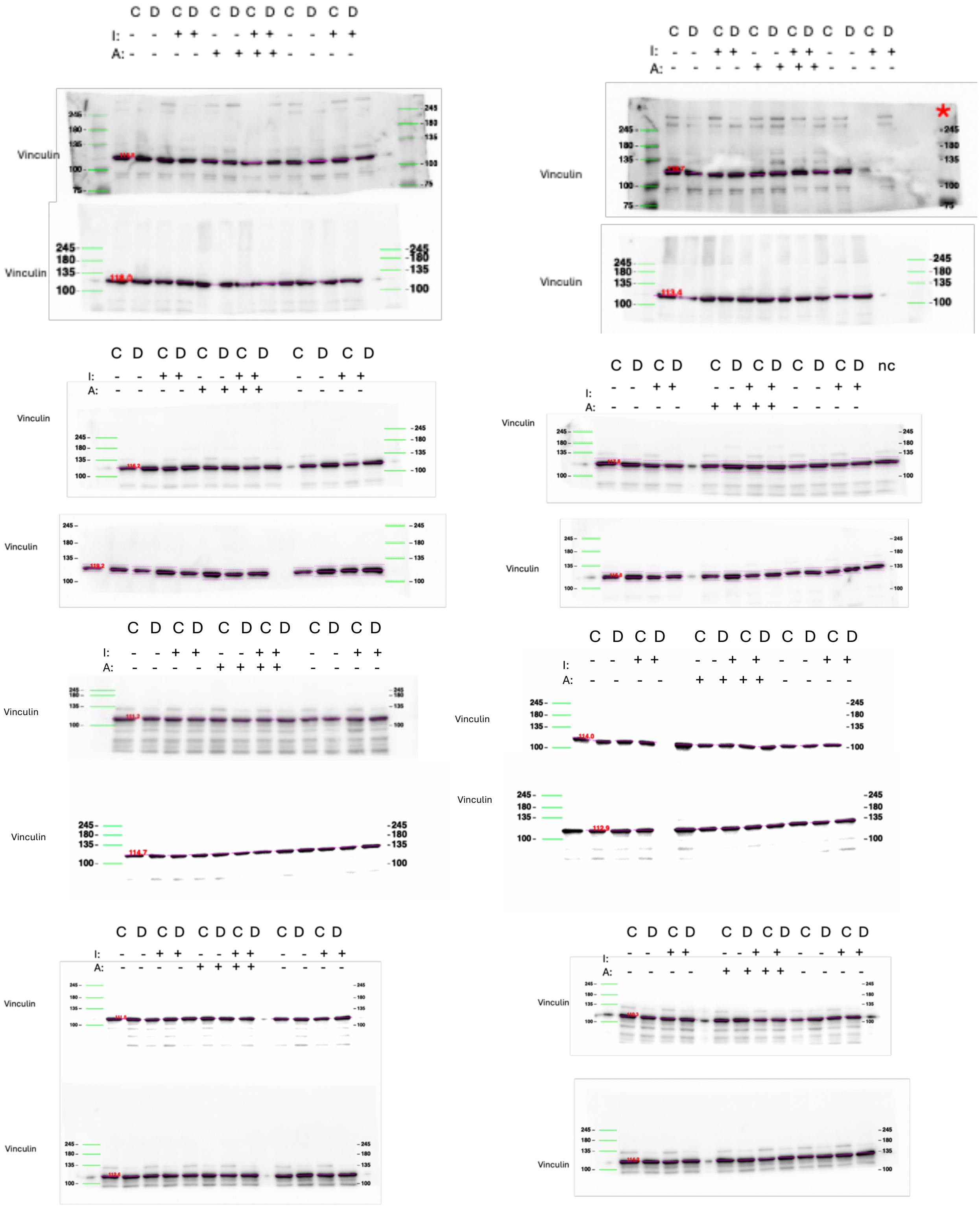

C D C D C D C D C D C D
I: - - + + - - + + - - + +
A: - - - - + + + + - - - -
Vinculin
245
180
135
100
75
C D C D C D C D C D C D
I: - - + + - - + + - - + +
A: - - - - + + + + - - - -
Vinculin
*
C D C D C D C D C D C D
I: - - + + - - + + - - + +
A: - - - - + + + + - - - -
Vinculin
C D C D C D C D C D C D nc
I: - - + + - - + + - - + +
A: - - - - + + + + - - - -
Vinculin
C D C D C D C D C D C D
I: - - + + - - + + - - + +
A: - - - - + + + + - - - -
Vinculin
C D C D C D C D C D C D
I: - - + + - - + + - - + +
A: - - - - + + + + - - - -
Vinculin
C D C D C D C D C D C D
I: - - + + - - + + - - + +
A: - - - - + + + + - - - -
Vinculin
C D C D C D C D C D C D
I: - - + + - - + + - - + +
A: - - - - + + + + - - - -
Vinculin

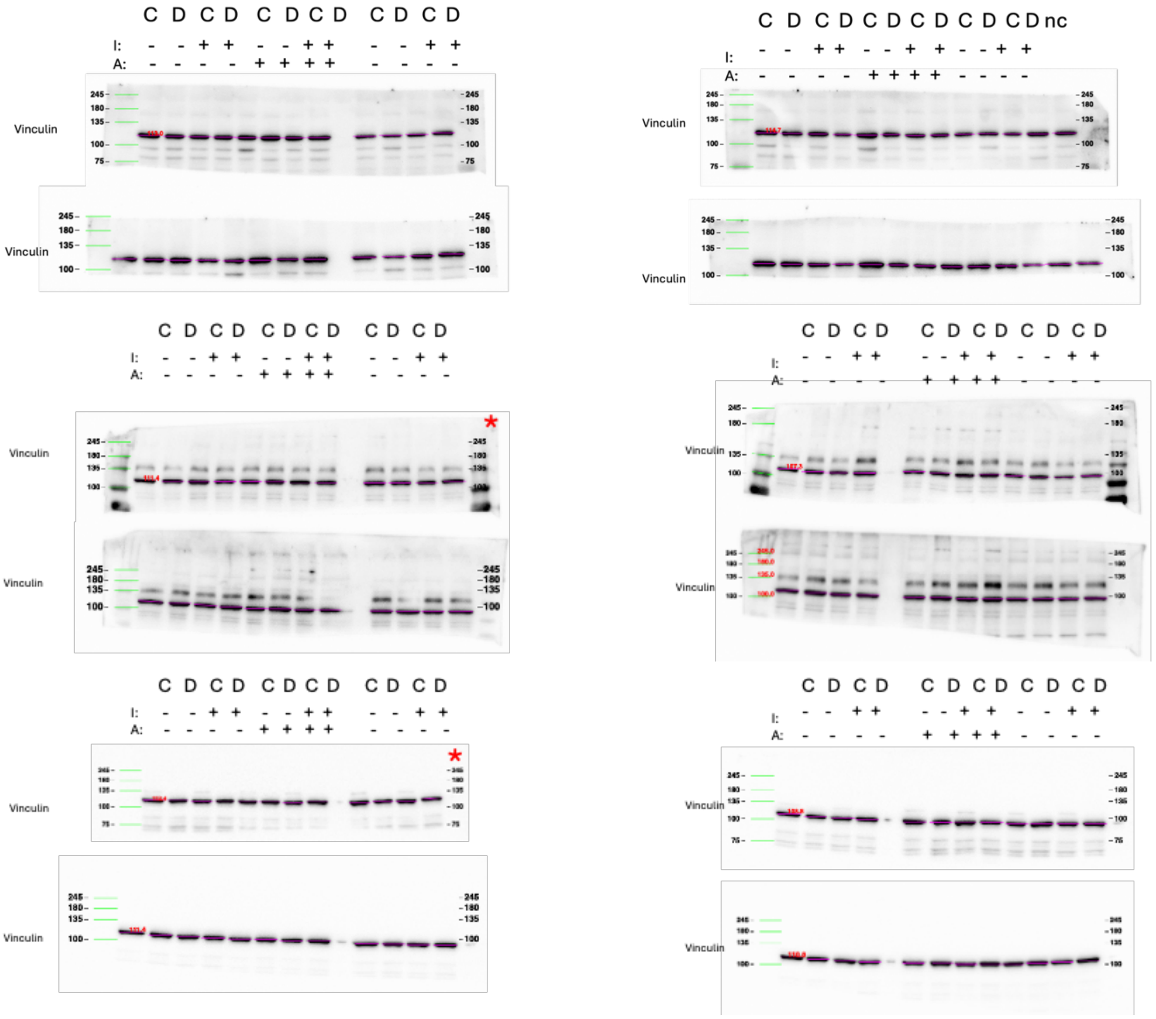

Supplemental Figure 16. Uncropped total vinculin myotube western blots for Figure 5A, 6A, 7A. Expected banding at ~124 kDa. Western blot banding found between ~110-120 kDa (Pink lines). Red asterisks indicate the blot used for representative blot. “C” represents Control myotubes and “D” represents DM1 myotubes. “I” represents insulin treatment and “A” represents AICAR treatment. ”nc” represents normalizing control.

**Supplemental Table 1. List of Antibodies Used in Western Blotting**.

| Primary Antibody | Company, Catalogue Number, RRID | Dilution | Secondary Antibody | Company, Catalogue Number, RRID | Dilution |
|---|---|---|---|---|---|
| Mouse anti-Vinculin | Santa Cruz, 73614, RRID:AB_2941767 | 1:2000 | Goat anti-mouse horse radish peroxidase (HRP) | Thermo Fisher Scientific, 31430, RRID:AB_228307 | 1:5000 |
| Rabbit anti-ACC | Cell Signaling, 3662S, RRID:AB_2219400 | 1:1000 | Goat anti-rabbit HRP | Thermo Fisher Scientific, 31460, RRID:AB_228341 | 1:5000 |
| Rabbit anti-pACC (Ser79) | Cell Signaling, 3661, RRID:AB_330337 | 1:500 | | | |
| Rabbit anti-AS160 | Cell Signaling, 2447S, RRID:AB_2199376 | 1:1000 | | | |
| Rabbit anti- | Cell Signaling, 4288S, RRID:AB_10545274 | 1:1000 | | | |

| | | |
|---|---|---|
| pAS160 (Thr642) | | |
| Rabbit anti-PGC-1α | Thermo Fisher Scientific, PA5-72948, RRID:AB_2718802 | 1:500 |
| Rabbit anti-AKT | Cell Signaling, 9272, RRID:AB_329827 | 1:1000 |
| Rabbit anti-pAKT (Ser473) | Cell Signaling, 9271S, RRID:AB_329825 | 1:1000 |
| Rabbit anti-AMPK | Cell Signaling, 2532S, RRID:AB_330331 | 1:1000 |
| Rabbit anti-pAMPK (Thr172) | Cell Signaling, 2531S, RRID:AB_330330 | 1:500 |
| Rabbit anti-GLUT4 | Abcam, ab33780, RRID:AB_2191441 | 1:1000 |

Supplemental file 2. List of molecular targets identified in sedentary HSA-LR versus sedentary WT condition

See excel file *Molecular Targets sed vs wt*

Supplemental file 3. List of molecular targets identified in exercised HSA-LR versus sedentary HSA-LR condition

See excel file *Molecular Targets ex vs sed*

Supplemental file 4. List of all canonical pathways identified in sedentary HSA-LR versus sedentary WT condition

See excel file 4.*canonical pathways sed vs wt*

Supplemental file 5. List of all canonical pathways identified in exercised HSA-LR versus sedentary HSA-LR condition

See excel file 5.*canonical pathways ex vs sed*

Supplemental file 6. List of molecular targets identified in PI3K-AKT canonical pathway in HSA-LR versus sedentary WT condition

See excel file 6.*pi3k akt sed vs wt*

Supplemental file 7. List of molecular targets identified in PI3K-AKT canonical pathway in exercised HSA-LR versus sedentary HSA-LR condition

See excel file 7.*pi3k akt ex vs sed*

Supplemental file 8. List of molecular targets identified in SLC2A4 (GLUT4) translocation canonical pathway in HSA-LR versus sedentary WT condition

See excel file 8.*glut4 translocation sed vs wt*

Supplemental 9. List of molecular targets identified in SLC2A4 (GLUT4) translocation in exercised HSA-LR versus sedentary HSA-LR condition

See excel file 9.*pi3k glut4 translocation ex vs sed*

Supplemental 10. List of upstream regulators identified in HSA-LR versus sedentary WT condition

See excel file 10.*upstream regulators sed vs wt*

Supplemental 11. List of upstream regulators identified in exercised HSA-LR versus sedentary HSA-LR condition

See excel file 11.*upstream regulators ex vs sed*
